# Continuous integration of heading and goal directions guides steering

**DOI:** 10.1101/2024.10.24.620060

**Authors:** Anthony M Crown, Annie H Wu, Lindsey Hofflander, Gilad Barnea

## Abstract

Navigating animals must integrate a diverse array of sensory cues into a single locomotor decision. Insects perform intricate navigational feats using a brain region termed the central complex in which an animal’s heading direction is transformed through several layers of circuitry to elicit goal-directed locomotion. These transformations occur mostly in the fan-shaped body (FB), a major locus of multi-sensory integration in the central complex. Key aspects of these sensorimotor computations have been extensively characterized by functional studies, leveraging the genetic tools available in the fruit fly. However, our understanding of how neuronal activity in the FB dictates locomotor behaviors during navigation remains enigmatic. Here, we manipulate the activity of two key neuronal populations that input into the FB–the PFN_a_ and PFN_d_ neurons–used to encode the direction of two complex navigational cues: wind plumes and optic flow, respectively. We find that flies presented with unidirectional optic flow steer along curved walking trajectories, but silencing PFN_d_ neurons abolishes this curvature. We next use optogenetic activation to introduce a fictive heading signal in the PFNs to establish the causal relationship between their activity and steering behavior. Our studies reveal that the central complex guides locomotion by summing the PFN-borne directional signals and shifting movement trajectories left or right accordingly. Based on these results, we propose a model of central complex-mediated locomotion wherein the fly achieves fine-grained control of sensory-guided steering by continuously integrating its heading and goal directions over time.

## Main

Insects perform complex navigational tasks with relatively simple nervous systems. These tasks vary in range and complexity, from path integration in foraging ants over hundreds of meters^1^ to the seasonal migration of monarch butterflies over thousands of kilometers^2^. Despite this vast range in navigational capabilities, a brain region conserved across insect species–the central complex–is thought to underlie these behaviors^3–5^. The central complex consists of four main compartments (Fig. 1a) that communicate via several populations of columnar neurons, the architecture and synaptic connectivity of which have been delineated in the fruit fly, *Drosophila melanogaster*^6,7^. One such compartment, the ellipsoid body (EB), intrinsically generates a representation of the fly’s world-centric, also known as allocentric, orientation in space. This representation takes the form of a neuronal activity “bump” within a circular arrangement of columnar neurons termed the EPGs that is yoked to the fly’s heading when landmarks are present (Fig. 1b)^8–12^. When flies perform navigational tasks that require a stable landmark, the EB activity bump is recruited^11,13^. In parallel, neural projections to the noduli (NO) are thought to encode body-centric, also known as egocentric, left-right sensory signals^14–16^. A population of neurons termed PFNs receives information from both pathways^6^ and conveys this information to the FB (Fig. 1b, c), a brain center where various sensory cues^17–19^ and aspects of the animal’s internal state^19,20^ are represented. The PFNs incorporate the egocentric sensory signals to transpose the heading signal and construct allocentric vector representations of these dynamic sensory cues^14,16^. Different PFN subpopulations perform this vector transposition for different complex navigational cues, such as wind plumes^15^ and optic flow^14,16^. The processing of these vector codes of sensory information culminates in the generation of a goal signal, which is represented as an activity bump in the FB. This goal signal, also expressed in allocentric coordinates, is used to guide locomotion during navigation^21,22^. Thus, the anatomical and functional properties of PFNs render them the likely origin of the fly’s goal-oriented steering behavior. Further, these neurons likely act as a key circuit node for transforming navigational cues into locomotion. Hence, we sought to determine the contribution of neuronal activity in the PFNs to navigational behaviors through thermo- and optogenetic manipulation of genetically defined subpopulations of PFNs.

**Figure 1.**
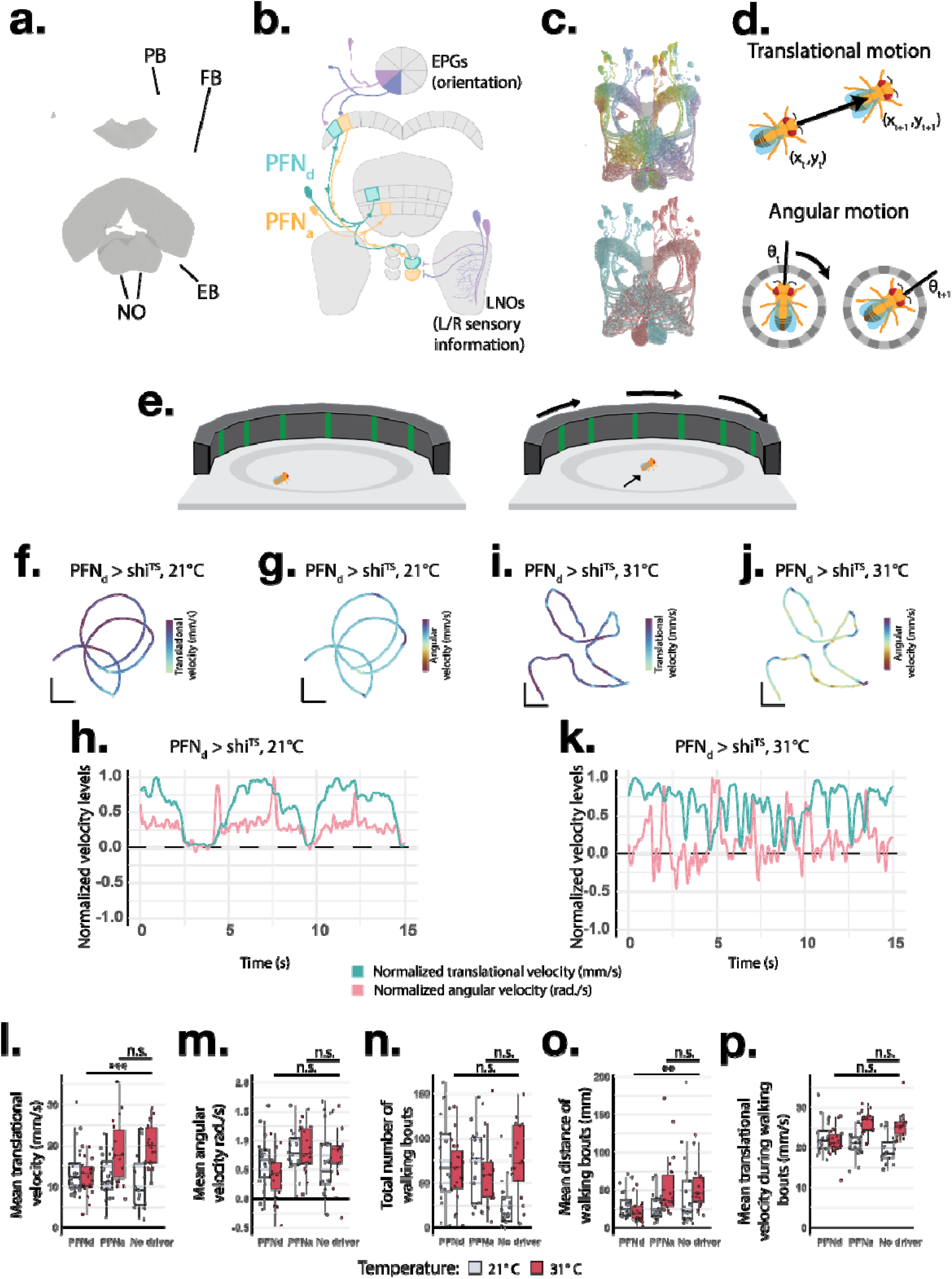
PFNs are necessary for controlling curvature of walking trajectories. **(a)** EM reconstructions of the main compartments of the central complex. **(b)** Schematic representation of the flow of information to the PFNs. EPG neurons (two shades of purple, above), encode the fly’s heading direction. Axons from an EPG neuron innervate a column of the PB. LNO neurons (two shades of purple, below) encode various left-vs-right sensory signals. Each LNO neuron innervates either a left or right nodulus. LNO neurons that encode different sensory signals innervate different noduli. Single PFN_a_ neurons (orange) and PFN_d_ neurons (blue) receive input from a single PB column and a single nodulus. **(c)** EM reconstructions of PFN neurons colored by either their FB column position (above) or the noduli from which they receive input (below). **(d)** Parameters used for quantifying translational (above) and angular (below) locomotion. **(e)** Cartoon representation of the behavioral paradigm. Flies are presented with unidirectional optic flow, which is locked in closed loop with their translational motion. **(f-k)** Representative walking trajectories from a unidirectional optic flow assay. Flies express the thermogenetic silencer Shibire^TS^ in PFN_d_ neurons. (f-k) depict 15s of walking at the permissive 21°C (f-h) and restrictive 31°C (i-k) temperatures. Trajectories are colored to depict either translational (f, i) or angular (g, j) velocity values. (h, k) depict translational velocity (cyan) and angular velocity (red) values over the course of the 15s walking bout. Velocity values were normalized to enable comparisons of how the two values vary over time. **(l-p)** Boxplots depicting changes in mean translational (l) and angular (m) velocity values, total number of walking bouts (n), average distance of walking bouts (o), and average translational velocity values during walking bouts (p) for 21°C and 31°C trials. Positive angular velocity values in (g, h, j, k, m) are towards the direction of the stimulus. One-way ANOVA with post-hoc Tukey’s multiple comparison test was used for statistical testing (* = p < 0.05, ** = p < 0.01, *** = p < 0.001). Scale bars in f, g = 10mm, i, j = 20mm.

### PFNs control body orientation during forward walking bouts

We first wished to devise a behavioral paradigm in which the PFN circuitry would be recruited. PFNs transform sensory signals from ego-to allocentric coordinate systems^14,16^. Different PFN populations perform this transformation for distinct sensory cues: PFN_a_ neurons represent wind plumes, and PFN_d_ neurons represent optic flow^14–17^. Both wind plumes and optic flow are complex cues that inherently convey relevant directional information to the navigating fly. Therefore, the fly must incorporate information from both when computing its movement decision. PFN_a_ and PFN_d_ neurons, however, target nonoverlapping regions of the FB (Fig. 1b) and input onto divergent circuits^6^. We hypothesized that PFN_a_ and PFN_d_ neurons perform parallel functions in navigational behaviors. We hence designed a behavioral paradigm that would rely on the activity in one subpopulation of PFNs (PFN_a_ or PFN_d_ neurons) but not the other. We decided to focus on the characterization of how optic flow contributes to navigational behaviors because of the simpler nature of the signal and the ability to precisely manipulate it compared to the dynamic and unpredictable nature of turbulent wind plumes.

When a landmark is present and other sensory cues are absent, flies display a behavior termed menotaxis in which they maintain a stable heading in an arbitrary goal direction relative to the landmark^11,23,24^. In addition, it incorporates self-motion cues to ensure that its movement is aligned with this goal direction. We hypothesized that optic flow constitutes one of these self-motion cues and that the fly uses PFN_d_-borne representations of optic flow to maintain a stable heading trajectory. We, therefore, sought to characterize how flies respond behaviorally to manipulations that misalign the optic flow and their heading direction as well as how activity in PFN_d_ neurons, in turn, affects these responses.

We thus designed a behavioral paradigm in which flies were presented with unidirectional optic flow as they move through space. To this end, we placed the flies in a circular chamber^25^ that is surrounded by green LED arrays programmed to display a series of vertical bars (Extended Data Fig. 1). By rotating the positions of the vertical bars around the circumference of the arena in a clockwise or counterclockwise fashion, the LED arrays produced an optical illusion of rotational movement. We tethered this visual stimulus to the fly’s movement by rotating it only when the fly moves so that the stimulus would better mimic optic flow cues indicating self-motion (Fig. 1e). We recorded videos of the behavioral responses of the flies to this stimulus paradigm and performed automatic kinematic tracking^26^ to quantify the results.

During locomotion, a fly must continuously update its body position (translational motion) and orientation (angular motion) in space (Fig 1d). We quantified both variables in the kinematic data and used these metrics to determine how the flies responded to our stimulus paradigm. We first presented control flies with unidirectional optic flow during bouts of forward walking. We found that these flies exhibited circular movement trajectories when close to the center of the arena (Fig. 1f-h, Extended Data Fig. 2a-c). Further, angular velocity values shifted significantly in the direction of the rotating stimulus and coincided with bouts of translational velocity (Extended Data Fig. 2d, e). While eliciting these trajectories, flies would maintain a stable angular velocity during discrete bouts of continuous movement that last upwards of 10s (Extended Data Fig. 2c). We termed these epochs of circular movement trajectories “reorientation bouts.” We then sought to understand how the brain coordinates translational and angular motion to produce reorientation bouts.

We hypothesized that flies elicit reorientation bouts by updating their internal goal coordinate to align with visual feedback that acts as a self-motion cue. We, therefore, wished to examine whether these reorientation bouts relied on neuronal activity in core components of the central complex. We thus thermogenetically silenced targeted populations of central complex columnar neurons in walking flies while subjecting them to our behavioral paradigm. To achieve that, we used driver lines that narrowly target individual neuronal subpopulations in the central complex^7^ to express shibire^TS^, an allele that reversibly blocks neurotransmission at temperatures greater than 28°C. We first turned to the EPG neurons, which encode the heading direction of the fly and act as a “master compass” for the central complex^11,24,27,28^. Flies of this cohort spent significantly less time in the center of the arena where curved walking trajectories are typically observed (Extended Data Fig. 3). Instead, these flies were usually located at the edge of the arena chasing the stimulus at a range where only one bar would be visible to them. This behavior is reminiscent of bar fixation, a behavior not thought to be controlled by the central complex circuitry. This result is thus aligned with previous observations that flies resort to more reflexive orientation behaviors in the absence of EPG neurons^11,24^. Since reorientation bouts are only observed in the center of the arena, we observed no such behavior when the EPGs were silenced. We, therefore, conclude that the central complex circuitry is engaged in our behavioral paradigm and that neuronal activity in the EPGs is likely used to produce the curved reorientation bouts.

The EPG activity bump, which represents the fly’s heading direction, is duplicated across the protocerebral bridge (PB), where it is inherited by neurons termed PFNs, the next layer of the central complex columnar circuit. PFNs incorporate asymmetric sensory signals from the NO to modulate the amplitude of the left and right activity bumps. The sum of the left and right PFN activity bumps produces a new activity bump in the next layer of circuitry. This new activity bump represents the direction of a particular sensory cue that is now transformed into allocentric coordinates^14,16^. These transformations ultimately generate a “goal direction” activity bump in the FB, which is compared to the heading direction represented by the EPG activity bump in the EB. The relative positions of the EB heading direction, and the FB goal direction are thought to drive angular movement during epochs of goal-directed locomotion^21,22^. We, therefore, predict that silencing PFNs would result in constitutive alignment of the heading and goal signals. Thus, it would prevent the fly from eliciting movements to align the two coordinates. To test this prediction, we used shibire^TS^ to selectively silence two populations of PFNs previously implicated in sensory guided forward motion–the PFN_a_^15,17^ and PFN_d_^14,16^ subtypes–while the flies perform the angular motion assay.

For flies performing the goal-directed walking assay at 21°C–a permissive temperature at which shibire^TS^ allows neurotransmission–angular motion was preserved, and the flies performed reorientation bouts across experimental groups (Fig. 1f-h, l-m). By contrast, flies bearing shibire^TS^ in PFN_d_ neurons that underwent the behavioral paradigm at 31°C–a restrictive temperature at which shibire^TS^ blocks neurotransmission–exhibited a “stop-and-turn” phenotype; their walking trajectories were straightened, and they only turned between bouts of forward walking (Fig. 1i-k). Hence, at the restrictive temperature, the values of translational and angular velocity were no longer coincidental (Fig. 1k). Over the duration of each trial, the translational velocity values in flies with silenced PFN_d_ neurons were significantly reduced, indicating deficits in forward motion (Fig. 1l). Similarly, the angular velocity values in these flies were shifted towards zero, albeit slightly (Fig. 1m). Notably, PFN_a_ neurons were dispensable for this behavior (Fig. 1l, m), indicating that the PFN_a_ and PFN_d_ subtypes perform specialized roles in navigational behaviors, which is in line with our predictions.

We observed that flies in the behavioral paradigm tended to walk forward in short bouts of translational velocity followed by periods of resting. In a five-minute experiment, animals with silenced PFN_d_ neurons walked overall shorter distances than control animals, as represented by the translational velocity values (Fig. 1l). This decrease in translational velocity could reflect a reduction in overall walking speeds. Alternatively, it could result from shorter bouts of activity at comparable speeds to control flies, resulting in a lower average walking speed overall. To differentiate between these possibilities, we decomposed our kinematic tracking data for each fly into the discrete walking bouts elicited within each trial. This analysis enabled us to determine the number of walking bouts, average distance travelled in each bout, and average translational velocity for each bout. When comparing these metrics to the control animals, we found that silencing PFN_d_ neurons led to no change in the number of walking bouts (Fig. 1m), but to a significant decrease in the overall distance travelled in each bout (Fig. 1n). The translational velocity values during these bouts of movement were comparable to those observed in control animals (Fig. 1o). Thus, this analysis indicates that the decrease in translational velocity observed when silencing PFN_d_ neurons is due to shorter bouts of movement at comparable velocities to those of controls. These observations are consistent with a role for PFN_d_ neurons in eliciting continuous bouts of forward movement with curved walking trajectories. We conclude that PFNs are necessary for instructing locomotion by driving bouts of forward walking that are curved towards the left or right direction.

### Additive effects of parallel PFN pathways on locomotion

It is noteworthy that silencing PFN_d_ neurons resulted in angular motion that was diminished but not altogether abolished. Therefore, it is possible that PFNs constitute one of multiple parallel steering systems and that the other systems are not perturbed by our manipulations. Such an organization would enable fine-grained control of body orientation during movement. We hypothesized that neuronal activity in PFNs is sufficient to elicit changes in the fly’s locomotor behavior. Should this indeed be the case, we would predict that exogenous activation of PFNs would elicit a shift in the fly’s ongoing heading direction in the absence of any navigational cues. To test this prediction, we optogenetically activated PFNs during locomotion and quantified changes in the fly’s heading. We expressed the light-gated cation channel CsChrimson in either PFN_a_ or PFN_d_ neurons using our selective driver lines. However, our thermogenetic silencing experiments revealed that PFN_a_ neurons were dispensable in the optic flow assay. This result may indicate that the two subtypes of PFNs are functionally subdivided and hence constitute parallel circuits. The distinct cues represented by PFN_a_ and PFN_d_ neurons both affect navigational behaviors. Further, the ultimate heading direction should be coherent and represent the integration of both cues. We thus hypothesized that the parallel PFN circuits sum their signals to produce a single, unified heading direction. To test this hypothesis, we employed an additional driver line that targets both PFN_a_ and PFN_d_ subtypes (PFN_a+d_) to simultaneously activate both populations. We activated the PFNs in freely walking flies by subjecting them to an optogenetic stimulus paradigm during which a red light illuminates the behavioral arena for 20s with 20s resting periods before and after each stimulation. Each fly underwent three total stimulation periods while in an otherwise dark chamber (Fig 2a). Because red light is not detected by the fly visual system^29^, this experimental design enables us to profile changes in the fly’s locomotion without introducing a goal stimulus through visual cues.

**Figure 2.**
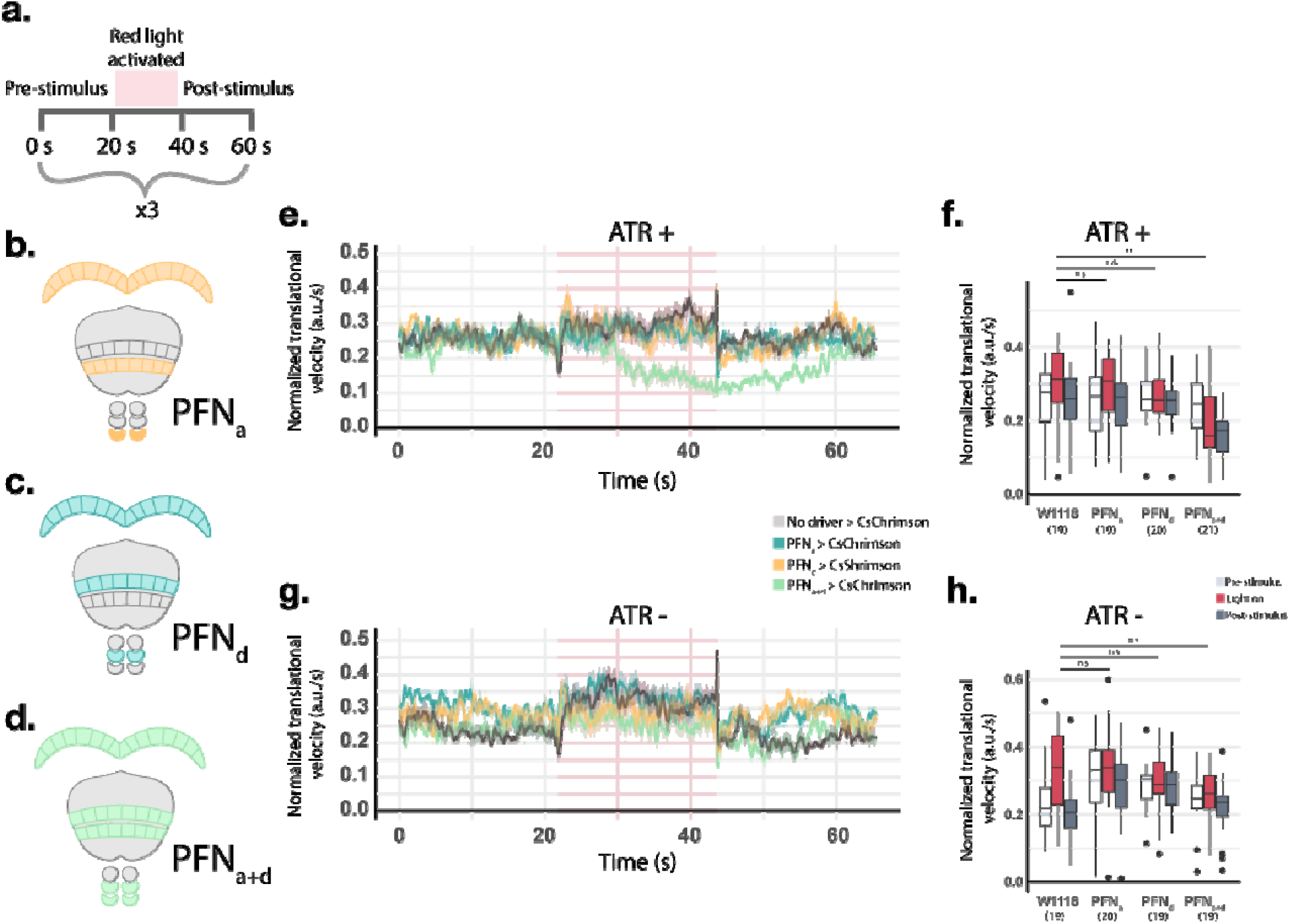
Simultaneous activation of PFN_a_ and PFN_d_ neurons suppresses locomotion. **(a)** Paradigm for optogenetic activation experiments. Flies are exposed to 20s of red light, with 20s rest periods before and after each stimulus. Each fly is exposed to three stimulus bouts total. **(b-d)** Cartoon depiction of the cell-types targeted by each driver line. Driver lines that target either PFN_a_ neurons individually (b, blue), PFN_d_ neurons individually (c, yellow), or both PFN_a_ and PFN_d_ populations (d, green), were employed. **(e)** line plot depicting averaged translational velocity values (±s.e.) for each stimulus bout for the various genotypes. Flies were raised on diet supplemented with all-trans retinal (ATR+), the necessary cofactor for CsChrimson. Red box indicates time interval when the optogenetic stimulus was delivered. See (f) for the numbers of trials (N) that were averaged in each group. **(f)** Boxplot of mean translational velocity values during each optogenetic stimulus bout for the various genotypes in the ATR+ condition. Values are shown for the optogenetic stimulus period as well as for pre- and post-stimulus periods. **(g)** and **(h)** correspond to (e) and (f) respectively, but for flies that were raised on diets without ATR. In e-h, translational velocity values for each fly were normalized such that the maximum for each fly equals 1 to account for variability in flies’ walking speeds. One-way ANOVA2 with post hoc Tukey’s multiple comparison test was used for statistical testing. (** = p < 0.01)

Activation of PFN_a_ or PFN_d_ neurons alone elicited no observable changes in translational (Fig. 2e-h) or angular (Extended Data Fig. 4a, b) velocity. However, simultaneous activation of both populations via our PFN_a+d_ driver resulted in an overall decrease in locomotion. Interestingly, in some cases, optogenetic stimulation of the PFN_a+d_ neurons led flies to stop moving altogether and completely freeze during the stimulation period (Extended Data Fig. 5). Plotting the averaged translational velocity values of flies during each stimulation bout reveals that translational motion continually decreases when PFN_a+d_ neurons are activated and slowly recovers to baseline levels at the end of the post-stimulus period (Fig. 2e). Thus, this line of investigation reveals that transient, simultaneous activation of both PFN_a_ and PFN_d_ populations results in a reversible cessation of locomotion.

### Asymmetric activation of PFNs biases heading direction

We hypothesized that PFN_a+d_-mediated suppression of locomotion is due to a summation of PFN_a_ and PFN_d_ heading signals in the FB. We further hypothesized that such a summation mechanism should exist to integrate the parallel sensory pathways into a single left/right motor command because the two PFN populations encode different classes of navigational cues^14–17^. However, since in our previous line of investigation we analyzed the contributions of the PFNs to locomotion by activating entire populations of PFN subtypes, we could not test this hypothesis. The general manipulations of entire PFN populations obscure the natural dynamics of these circuits because neuronal activity in the PFNs typically takes the form of a sinusoidal bump, the peak of which is localized in a particular column^9,14,16^. Therefore, we reasoned that uncovering a more nuanced relationship between neuronal activity in the PFNs and locomotion would require a more selective targeting of the activated PFN population. Such targeted manipulation of a subpopulation of PFNs would better recapitulate their functional dynamics, and thus, more accurately mimic a specific directional signal. Further, a more targeted analysis of PFNs would better elucidate the mechanism of how exogenous activation of PFN_a_ and PFN_d_ neurons controls steering behavior to allow for orientation. We thus employed a mosaic strategy to optogenetically activate sparse and stochastically selected subsets of the PFNs.

We stimulated sparse and stochastically selected subpopulations of the neurons targeted by the PFN_a+d_ driver line via SPARC^30^ and employed the same optogenetic paradigm we used to activate all PFNs of a particular subtype. We stimulated these sparse PFN subsets in the absence of any navigational cues and tracked the locomotor behaviors of the flies. We then employed a *post-hoc* dissection and immunohistochemical analysis to determine which PFN neurons were activated in each fly.

Sparse activation of PFNs led to observable shifts in angular velocity, typically following the onset of delivery of the optogenetic stimulus (Fig. 3e, f, black arrowhead). We investigated whether the direction and magnitude of these shifts in heading direction could be predicted based on which PFNs were activated in each experiment. Because left/right sensory information is conveyed to the PFNs via asymmetric activity in the NO^14,15^, we tested whether asymmetric PFN activation elicits unilateral shifts in the fly’s heading direction. To achieve this, we first computed an index for asymmetric labeling of PFNs in our SPARC experiments by quantifying CsChrimson::tdTomato fluorescent signal in the corresponding left and right noduli. This analysis allows us to quantify the levels of asymmetric PFN activation and compare these values to the changes in heading elicited upon optogenetic stimulation. Because PFN_a_ and PFN_d_ neurons can be distinguished by the noduli from which they receive input^6^, this strategy allows us to additionally profile the relative contributions of the individual PFN subpopulations to changes in heading. We term the indices of PFN asymmetry ΔNO_a_ and ΔNO_d_ for PFN_a_ and PFN_d_ neurons, respectively (Fig. 3a).

**Figure 3.**
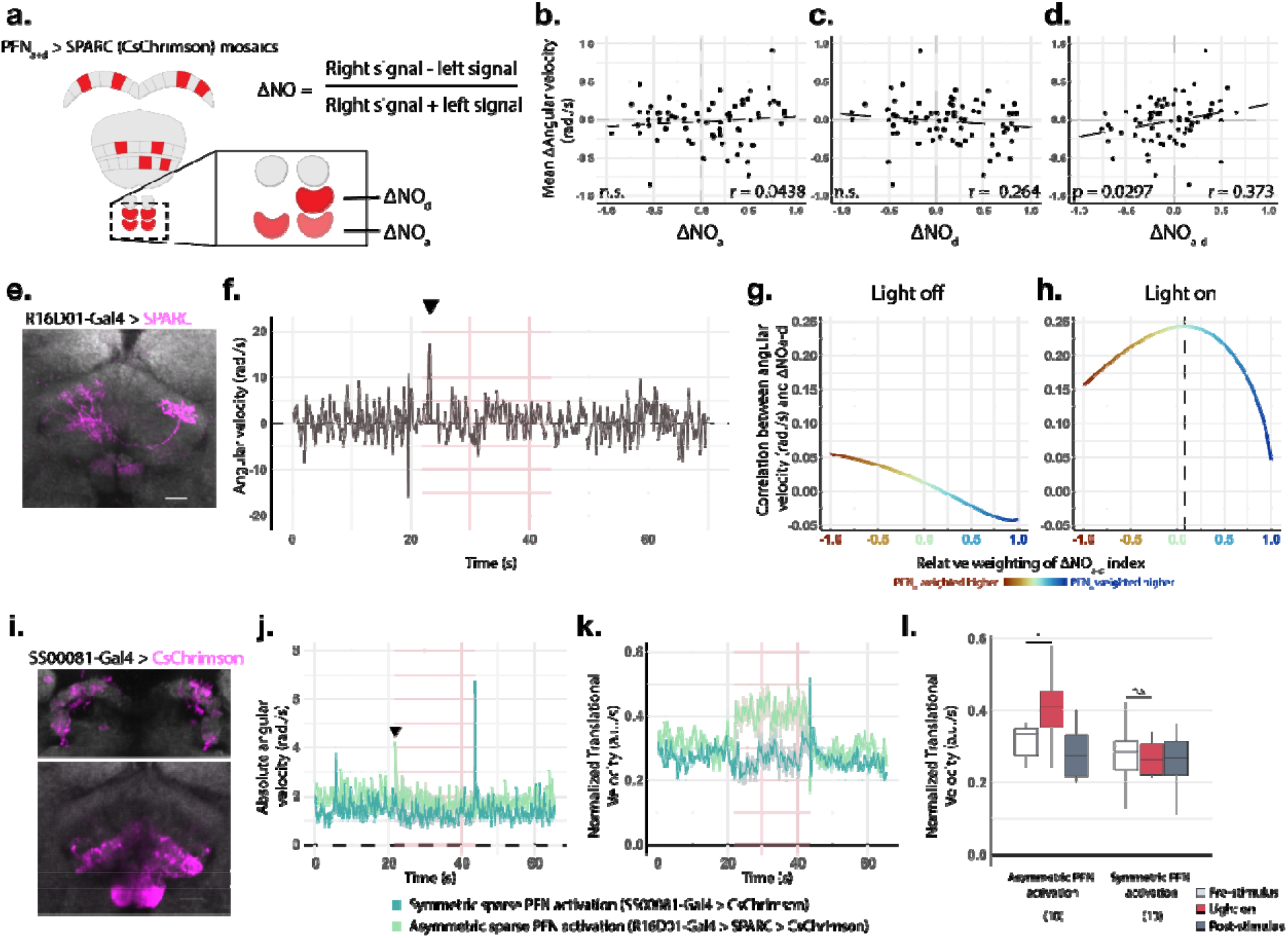
Asymmetric activation of PFNs shifts heading direction. **(a)** Schematic representation of how ΔNO indices of PFN asymmetry were calculated amongst PFN_a+d_ mosaic flies (see methods for details). **(b-d)** Scatterplot of PFN_a+d_ > SPARC experiments. Y-axis represents the change in mean angular velocity between time periods when the optogenetic stimulus was off versus when the optogenetic stimulus was on. X-axis depicts ΔNO indices for (b) PFN_a_ neurons (ΔNO_a_), (c) PFN_d_ neurons (ΔNO_d_), and (d) summed contributions of both populations (ΔNO_a-d_). r values represent the Pearson correlation coefficients. **(e)** Representative image of PFN_a+d_ > SPARC CsChrimson expression pattern (magenta) in the FB and NO. **(f)** Line plot depicting angular velocity values for the fly depicted in (e). One optogenetic stimulus bout is shown. **(g, h)** Line plot of relative contributions of PFN_a_ and PFN_d_ neurons to angular velocity values by computing weighted ΔNO_a-d_ indices. Y-axis shows the Pearson correlation between the ΔNO_a-d_ index and mean angular velocity values for PFN_a+d_ > SPARC CsChrimson trials. (g) and (h) depict the time intervals when the optogenetic stimulus was off and on respectively. X-axis and colors depict relative weighing of PFN_d_ neurons (red, −1) and PFN_a_ neurons (blue, +1). Dotted line in (h) denotes the maximum point. **(i)** Targeted expression pattern of CsChrimson (magenta) in the PB (upper panel) and the FB/NO (lower panel) via the sparse but symmetric PFN_a_ driver (SS00081-Gal4). **(j-k)** Line plots of averaged absolute angular velocity values (j) and normalized translational velocity values (k) when activating symmetric (SS00081-Gal4 > CsChrimson) or asymmetric (PFN_a+d_ > SPARC CsChrimson) sparse PFN subsets. Averaged values for ten trials shown. **(l)** Boxplot of mean normalized translational velocity values during the various optogenetic stimulation periods for trials depicted in (k). Scale bars in e, i = 10μm. One-way ANOVA with *post hoc* Tukey’s multiple comparison test was used for statistical testing in (i). (* = p < 0.05)

To quantify the contributions of asymmetric PFN activity to heading direction, we computed a Pearson correlation between the ΔNO_a_ or ΔNO_d_ indices and the mean changes in angular velocity upon optogenetic stimulation. This correlation was stronger for the ΔNO_d_ index than for the ΔNO_a_ index, indicating that PFN_d_ neurons elicited stronger changes in lateral movement (Fig. 3b, c). This result suggests that the activation of PFN_d_ neurons evokes stronger changes in heading direction than the activation of PFN_a_ neurons in this particular context.

Plotting the ΔNO_a_ or ΔNO_d_ indices against the mean changes in angular velocity for each experiment revealed the slopes of both trend lines (Fig, 3b, c). We inferred that the sign of these slopes, albeit statistically insignificant trends, indicates the direction of locomotor heading bias for each PFN population. The value of the ΔNO_a_ index increased as the measured changes in angular velocity became more positive (Fig. 3b). By contrast, the value of the ΔNO_d_ index increased as the measured changes in angular velocity became more negative (Fig. 3c). We then computed a third index for PFN asymmetry, ΔNO_a-d_, to quantify any interactions between PFN_a_ and PFN_d_ subtypes while accounting for the inferred sign of their contributions (see methods). Comparing the ΔNO_a-d_ index with the mean changes in angular velocity upon optogenetic activation revealed a statistically significant correlation (P=0.0297) between the two variables that was much stronger than the ΔNO_a_ or ΔNO_d_ indices alone (Fig. 3d).

To better assess the relative effects of PFN_a_ and PFN_d_ neurons on shifts in heading, we computed a weighted ΔNO_a-d_ index that varied from −1 to 1, with −1 indicating that only PFN_d_ neurons contributed to the index, 0 indicating that PFN_a_ and PFN_d_ neurons contributed equally, and 1 indicating that only PFN_a_ neurons contributed (see methods). During optogenetic stimulation, this weighted ΔNO_a-d_ index was most correlated with angular velocity when both populations were weighted about equally (ΔNO_a-d_ weight = 0.075) (Fig. 3g, h). We interpret these data as evidence for a summative effect between PFN_a_ and PFN_d_ neurons in controlling angular motion during movement. The observation that PFN_a_ and PFN_d_ neurons have opposing signs regarding their effects on movement may indicate that the two subsystems are configured in counterphase. Activation of all PFN_a_ and PFN_d_ neurons simultaneously may, therefore, lead to destructive interference between the two heading signals, ultimately suppressing locomotion, as we observed when activating all neurons contained in our PFN_a+d_ driver (Fig. 2e, f).

Our data indicate that the degree of symmetry of the sparse PFN population activated in a given fly affects its angular velocity. We would thus predict that activation of a sparse population of symmetric PFNs would not elicit changes in angular velocity. We tested this prediction by driving CsChrimson with a driver that targets a sparse but symmetric subset of PFN_a_ neurons (SS00081-Gal4) and subjected these flies to the optogenetic stimulation paradigm. We compared these trials to the same number of a randomly sampled subset of the PFN > SPARC CsChrimson trials. Remarkably, only in the PFN > SPARC experiments did we observe a spike in angular velocity at the onset of delivery of the optogenetic stimulus. This observation is consistent with a bout of reorientation upon asymmetric PFN activation. By contrast, SS00081 > CsChrimson flies exhibited a spike in angular velocity only upon offset of the opotogenetic stimulus (Fig. 3j). That symmetric PFN activation elicited reorientation upon stimulus offset could have indicated that angular motion was suppressed during optogenetic stimulation. However, we observed a similar spike in angular motion in a control experiment using SS00081 > CsChrimson animals raised on a diet lacking all-*trans* Retinal (ATR), a necessary cofactor for CsChrimson functionality (Extended Data Fig. 6). We, therefore, suspect that the offset spike in angular velocity values observed during symmetric PFN activation (SS00081 > CsChrimson flies) is an artifact of the optogenetic stimulus delivery. Intriguingly, only PFN > SPARC CsChrimon flies, but not SS00081 > CsChrimson flies, exhibited increased translational velocity during the optogenetic stimulus bout (Fig. 3k, l). Our interpretation of these observations is that indeed, asymmetric, but not symmetric, activation of PFNs elicits reorientation and forward motion.

### Antiphase relationship between PFN_a_ and PFN_d_ neurons predicted by the connectome

Asymmetric activation of either PFN_a_ or PFN_d_ neurons leads to a shift in angular motion, but the effects of the two populations are in opposite directions (Fig. 3b-d). These results could indicate that the two populations of neurons are arranged in an antiphase configuration. We, therefore, sought to examine whether the circuit connectivity of the FB would predict antiphasic relation between PFN_a_ and PFN_d_ neurons. To achieve that, we mapped the circuitry downstream of the two PFN populations using a recently completed connectome of a fly brain^31^. In our connectomic studies, we focused on the main postsynaptic targets of the PFN neurons, the hDelta interneurons. We, hence, sought to identify a pathway that would link PFNs via the hDelta interneurons to PFL3 neurons, a population of columnar neurons that translates goal signals in the FB into premotor steering commands^21,22^ (Fig. 4a).

**Figure 4.**
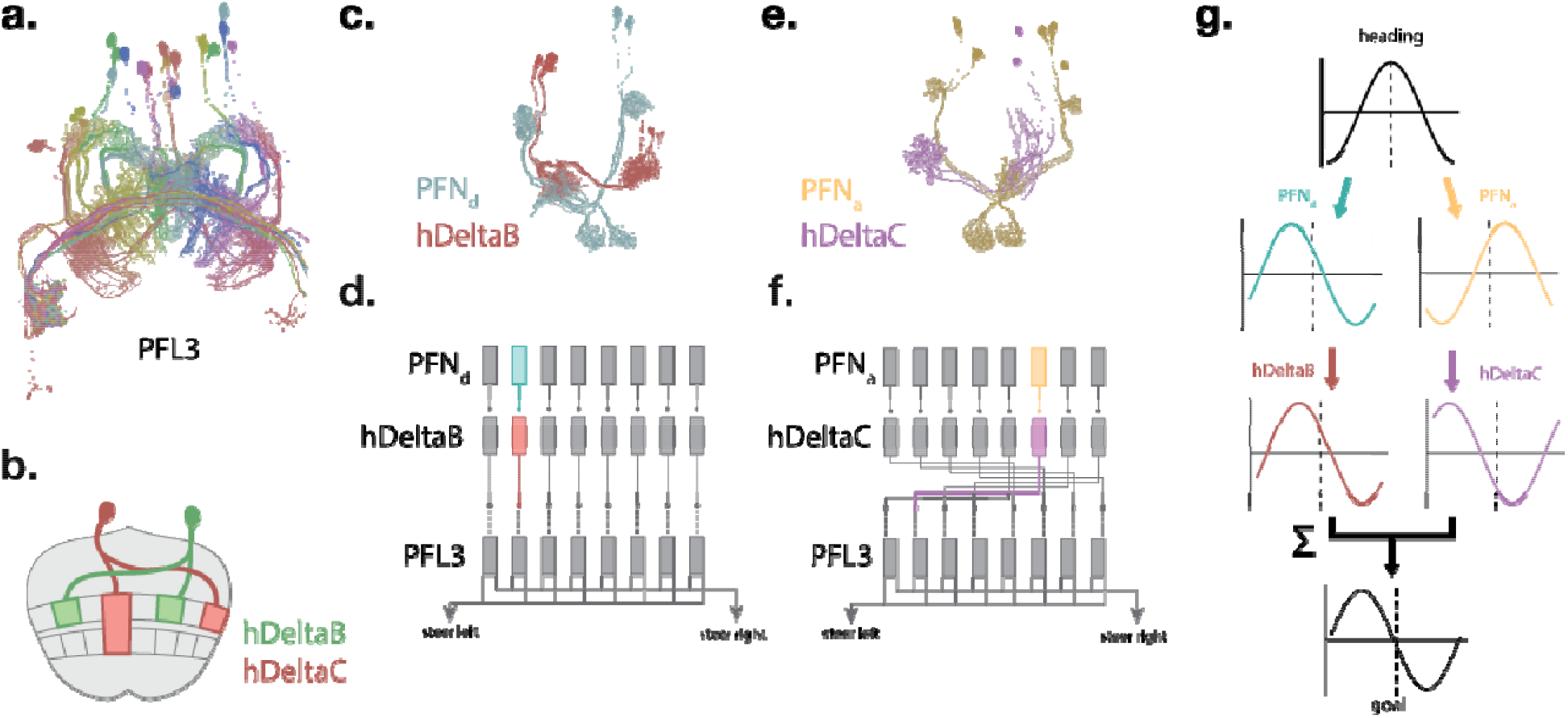
Mapping of PFN-born sensory signals onto central complex output neurons. **(a)** EM reconstruction of PFL3s, putative outputs of the central complex circuit, colored by FB column position. **(b)** Cartoon depiction of hDeltaB and hDeltaC neurons. Each hDelta neuron projects both ipsilateral and contralateral neurites. The ipsi- and contralateral neurites of a given hDelta neuron are offset by four FB columns. **(c-f)** EM reconstructions of the PFN neurons and hDelta neurons that are upstream of the PFL3s. For simplicity, only the populations that are upstream of the PFL3 neurons in the second FB column are shown. (c) EM reconstructions of the PFN_d_ neurons and hDeltaB neurons that are upstream of the PFL3s. (d) Cartoon depiction of the mapping of PFN_d_ neurons onto the PFL3s. Colors correspond to neurons depicted in (c). (e) Same as (c) but for PFN_a_ neurons and hDeltaC neurons. (f) Same as (d) but for PFN_a_ neurons. Colors correspond to neurons depicted in (e). Note that in (e, f) the depicted PFN_a_ neurons are offset from the second FB column and instead innervate the sixth column of the FB. **(g)** Example of the transformations that the heading signal, depicted as a sine wave, undergoes before forming a goal signal. PFN_d_ and PFN_a_ neurons offset the heading signal based on left-vs right sensory information. hDeltaB neurons inherit the PFN_d_ sensory signal and leave it untransformed. hDeltaC neurons inherit the PFN_a_ sensory signal and invert its phase. The sum of the hDeltaB and hDeltaC signals forms a goal signal that is relayed to the PFL3s for eliciting steering commands. The dotted line depicts the position of the heading signal.

Each hDelta neuron innervates one ipsilateral and one contralateral column of the FB with an offset of four columns. Given the phasic organization of the FB, this offset corresponds to an approximately 180° shift (Fig. 4b). This morphology of the hDelta neurons positions them as a potential mediator of the antiphasic relation between the PFN_a_ and PFN_d_ neurons, which is then ultimately inherited by the PFL3 neurons via additional intermediate neurons. Should this indeed be the case, a given column of PFL3 neurons would receive information from one PFN subtype (a or d) via the ipsilateral neurites of hDelta neurons and from the other PFN subtype via the contralateral neurites of other hDelta neurons. In this manner, the sinusoidal activity bump of one PFN population would be offset by 180° and, hence inverted, while the activity bump of the other PFN population would be unperturbed.

To determine whether the hDelta neurons indeed perform this signal inversion, we mapped their connectivity. Our analysis revealed that PFN_d_ neurons connect to PFL3 neurons via hDeltaB neurons (Fig. 4c, d), and, further, that hDeltaBs map onto PFL3s via their ipsilateral neurites (Fig. 4d). Thus, the phase of the PFN_d_-borne sensory signal is likely untransformed through this layer of the circuit. By contrast, the PFN_a_ neurons connect to PFL3 neurons via the contralateral neurites of the hDeltaC neurons (Fig. 4e, f). This anatomy indicates that the sensory-scaled representation of the navigational vector from PFN_a_ neurons, but not from PFN_d_ neurons, is inverted before being incorporated into the fly’s goal signal (Fig. 4g). Thus, the antiphase relationship that we observed in our optogenetic activation experiments is likely due to the anatomical substrates that we identified in the FB connectivity.

### A model for PFN-instructed steering in walking flies

Our results suggest a mechanism wherein the fly controls its steering maneuvers by comparing its internal heading signal to an external goal direction. This mechanism is supported by functional analysis of the FB^21,22^. Our results further suggest that asymmetric activation of PFN neurons transposes the fly’s goal relative to its current heading. Because silencing PFNs produced straightened walking trajectories, we hypothesized that the fly continuously compares its heading with its goal to determine its angular velocity at any given moment. Thus, when PFNs are silenced, the heading and goal signals are constantly aligned, and when PFNs are asymmetrically activated, the fly performs a corresponding turn to align its heading with its goal direction (Fig. 5a-c). These behaviors indicate that the fly compares its heading and goal orientations during movement, and that the integration of these two parameters relative to each other over time enables the fly to produce smoothly curved walking trajectories.

**Figure 5.**
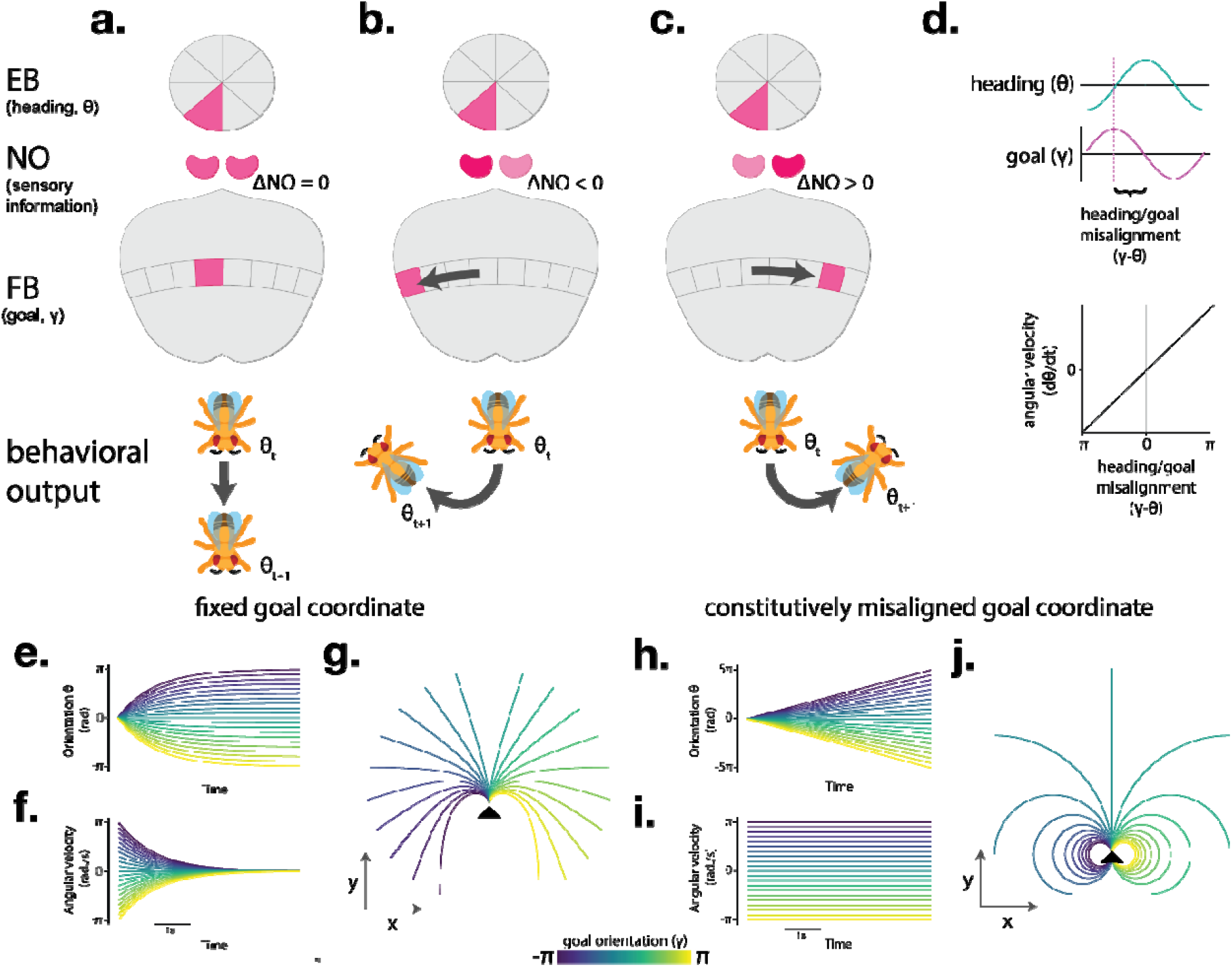
A model for central complex-mediated steering control. **(a-c)** Cartoon depictions of the relationship between heading signals, goal signals, and locomotion. From top to bottom: the heading signal in the EB, varying levels of asymmetric NO activation (depicted by color intensity), transposed goal signal in the FB, and hypothetical resultant walking trajectories (θ_t_ represents initial timepoint, θ_t+1_ represents end timepoint). Symmetric NO activation in (a) results in aligned heading and goal signals. Asymmetric activation in the NO as depicted in (b, c) transposes the goal signal and guides steering movements in the corresponding directions. **(d)** Parameters of model for central complex-mediated steering control. Angular velocity is directly proportional to the offset between heading and goal coordinates. **(e-g)** Modelling fly movement during orientation towards a fixed goal coordinate (dγ/dt = 0). Predicted orientation (d), angular velocity (e), and walking trajectories (f) are depicted. **(h-j)** Same as (e-g) but for modelling fly movement when heading and goal coordinates are constitutively misaligned at a fixed distance (dγ/dt = dθ/dt). Black arrowheads in (g, h) represent starting position for walking trajectories. Initial offsets between orientation and goal signals in (e-j) are displayed as γ values ranging from −π to π.

To gain insight into how these two parameters–heading and goal directions–are integrated continuously to produce movement with a smooth curvature, we sought to construct a mathematical model for this behavior. We thus constructed a series of ordinary differential equations wherein the relative positions of the fly’s heading (θ) and goal (γ) determine its angular velocity (dθ/dt). Recent studies have proposed that steering commands from the central complex are proportional in their magnitude to the degree of offset between θ and γ^21,22^. We hence modeled dθ/dt as proportional to (*i*.*e*., exhibiting a linear relationship with) the difference between θ and γ (Fig. 5d). We then sought to characterize the forward motion trajectories that such a relationship between θ and γ would produce when θ and γ are misaligned.

We defined γ as a static value (dγ/dt = 0), which produces a solvable system of ordinary differential equations. Varying the initial values for θ and γ in these solved equations demonstrates how angular velocity changes depending on the degree of offset between goal and heading signals. We thus initialized θ as equal to zero radians and varied the values of γ between −π and π radians to cover the full range of potential offset angles between heading and goal orientations. Plotting this modeled orientation θ over time showed smooth curves that were sharpest at the onset of the experiment before settling to straight lines as θ approached γ (Fig. 5e). Similarly, plotting the angular velocity dθ/dt over time revealed curves that were sharpest during the onset of the experiment but resolved to zero over time (Fig. 5f). This pattern of angular velocity values mirrors our results for optogenetic activation of PFNs, in which angular velocity values shifted most dramatically upon the onset of stimulation before returning to baseline levels (Fig. 3f, j).

We next modeled how these predicted angular velocity values would manifest as walking trajectories. Our behavioral data indicated that bouts of translational motion tend to be at a stable walking speed (Extended Data Fig. 2d). Thus, we calculated theoretical walking trajectories by computing the movements in the x and y directions given our modeled θ and assuming a constant walking speed. Our calculation produces smoothly curved walking trajectories that straighten once the flies are aligned to their respective γ values (Fig. 5g).

We next sought to define the parameters of the model such that they would reflect the fly’s heading and goal coordinates when presented with a unidirectional optic flow stimulus. We hypothesized that when flies are presented with optic flow that emulates rotational self-motion, their goal direction is continuously offset from the heading proportionally to the rotational speed of the optic flow. We modeled this by defining γ to rotate in concert with angular velocity (dγ/dt = dθ/dt). Such a configuration causes constitutive misalignment of θ and γ to various degrees according to the initial values of the two variables. As with our previous model, we initialized the value of θ to equal 0 and varied the values of γ between −π and π radians to cover the full range of potential offset angles between heading and goal orientations.

In this configuration of the model, θ changes linearly, and the rate of this change is proportional to the degree of offset between θ and γ (Fig. 5h). Hence, the angular velocity values are fixed and are manifested as straight lines when plotted over time (Fig, 5i), mimicking the stable angular velocity values we observed in reorientation bouts (Fig. 1h, Extended Data Fig. 2b, c). Similarly, computing theoretical walking trajectories from the modeled values reveals circular movement patterns (Fig. 5j) that are analogous to those observed in reorientation bouts (Fig. 1f, g, Extended Data Fig. 2a). We conclude that these modeling results indicate a possible strategy used by the fly brain to integrate goal and heading signals to control the curvature of forward walking bouts during navigational behaviors.

## Discussion

Surprisingly little is known about how neuronal activity in the FB drives locomotor behaviors. Our studies address this clear gap in knowledge by establishing a causal relationship between neuronal activity in the PFNs, a major input population to the FB, and steering movements. We demonstrate that thermogenetic silencing of PFNs resulted in the inability to elicit forward walking bouts with curved trajectories, which we interpret as deficits in steering control. However, optogenetic activation of PFNs was not wholly sufficient to elicit curved walking trajectories, such as those observed when descending command neurons for forward walking are optogenetically activated^32^. Our studies instead indicate that the steering commands to align the heading and goal signals are elicited only during the onset of asymmetric stimulation of PFNs. Beyond this onset period, shifts in angular motion during PFN stimulation are more consistent with biases in steering direction. PFNs may, therefore, function under natural conditions to maintain a stable heading during navigational behaviors by allowing the fly to smoothly adjust its movement trajectories in response to a dynamic influx of sensory cues. Such a role for the PFNs would, therefore, explain their necessity for curved walking bouts in our optic flow assay despite the apparent lack of an explicit goal coordinate in this behavioral paradigm.

Optomotor behaviors, such as those exhibited for gaze stabilization, are thought to be reflexive, *i*.*e*. reliant on simple sensorimotor transformations^33–35^. Therefore, our observation that the neural circuitry of the central complex is used to produce reorientation bouts in response to optic flow is intriguing. It is well established that PFN_d_ neurons encode information about the direction of the optic flow and contribute to building vectorial representations of the fly’s ongoing travelling direction^14,16^. Nevertheless, how the information that PFN_d_ neurons encode is used by the fly to elicit goal-directed behaviors remains unknown. Our observation that PFN_d_ neurons are necessary for the fly to elicit curved bouts of continuous movement to align with the direction of optic flow may indicate that the fly uses self-motion cues to maintain a stable goal coordinate. If this is indeed the case, optic flow signals in the central complex can be likened to an error correction mechanism, where heading, travelling, and goal directions are continuously compared to compute movement decisions.

Our optogenetic activation experiments revealed that the PFN_a_ and PFN_d_ subtypes are correlated and anticorrelated with angular motion, respectively. We interpret these data to indicate that the two populations are configured in antiphase, an interpretation that is supported by our connectomic analyses of the FB circuit. This antiphase relationship may explain why the simultaneous activation of PFN_a_ and PFN_d_ neurons resulted in decreased locomotion as the sum of two counterphase sine waves results in destructive interference. In the most extreme case, such an interference would manifest as a complete cessation of movement, as we sometimes observed.

During navigational behaviors, animals must integrate disparate sensory cues into a single movement decision. The brain must, therefore, contain a mechanism for comparing these cues and executing locomotor behaviors in accordance. One such mechanism would employ a “winner takes all” strategy, where the brain weighs the various sensory cues and selects only the most salient for its heading decision. An alternative would employ a “summation” strategy in which the brain incorporates all the various relevant sensory cues and computes its movement decision accordingly. We found that the parallel sensory signals in PFN_a_ and PFN_d_ subpopulations and their relationships with locomotor behavior were consistent with a summation strategy. Similar summation mechanisms that function within PFN subtypes responding to the same sensory modality have been shown^14,16^. However, no such mechanism has been described to perform an analogous transformation across parallel PFN subsystems responding to different modalities. Our studies thus lay the groundwork for future research to identify the nodes of convergence between the PFN_a_ and PFN_d_ pathways that mediate these summative properties.

It is noteworthy that our optogenetic activation experiments were designed to study the effects of exogenous neuronal activity in the central complex while minimizing any influence from external sensory cues. Flies are capable of complex navigational behaviors like path integration while relying on an entirely idiothetic sense of space^36–38^. However, some of the PFNs analyzed in this study are known to be negatively correlated with heading in the absence of visual cues and positively correlated with heading in the presence of visual feedback^9^. Additionally, the influence of wind-tracking PFNs on movement depends on the presence and valence of odorants in the environment^17^. We, therefore, expect our results to represent only a narrow range of the properties demonstrated by this circuitry under natural conditions. Finally, since our manipulations were performed in walking flies, whether the mechanisms we identified extend to similar directional maneuvers during flight is yet unknown. That said, the PFN circuitry is indeed engaged during flight^9,16^. Therefore, we expect that in airborne flies the PFNs perform similar computations to those described in this study.

The activity bumps in the central complex operate as vectorial representations of sensory information^14,16^. The topographical organization of the central complex has rendered it an attractive system for studying how neural circuits can transform these vector codes into navigational behavior outputs. Such characterization has led to the establishment of basic principles for how ensembles of neurons can perform fundamental mathematical operations like vector addition^14,16,21,22^ and inversion^39^. Our studies contribute to this growing body of knowledge by revealing how the fly brain may integrate the relative positions of various vector codes over time to guide movement. These basic principles could potentially extend to vertebrate systems where the animal may perform more complex navigation tasks. Vector codes are indeed ubiquitous in the mammalian brain^40^, including head direction-representing cells that are analogous to the EB-born heading signal^41–43^. Further, modelling studies predict that the cognitive maps of space in the mammalian hippocampus are constructed via vectorial representations of environmental boundaries and landmarks^44,45^. Understanding the neural connectivity motifs underlying the function of the *Drosophila* central complex may, therefore, provide a fundamental basis for understanding how the brain performs navigational tasks in diverse animal species.

## Materials and methods

### Fly genetics

All fly stocks were maintained at either 18°C or 21°C on standard cornmeal-molasses-agar media. Crosses and their progeny, unless otherwise stated, were kept at 25°C in a humidity-controlled incubator with a 12-hour light and 12-hour dark cycle. The fly lines that were used in this study were as follows: W1118 (BDSC #5905), UAS-shibire^TS 46^, SS02255-Gal4 (BDSC #75923), SS00078-Gal4 (BDSC #75854), R16D01-Gal4 (BDSC #48722), SS00090-Gal4 (BDSC #75849), SS54549-Gal4 (BDSC #86603), SS00081-Gal4 (BDSC #75848), UAS-CsChrimson::Venus (BDSC #55135), nSyb-IVS-PhiC31 (BDSC #84151), UAS-IVS-PhiC31 (BDSC #84154), UAS-SPARC2-S-CsChrimson::tdTomato (BDSC #84145), UAS-SPARC2-I-CsChrimson::tdTomato (BDSC #84144).

### Locomotion assay

All behavior experiments were performed in a temperature and humidity-controlled chamber. Unless otherwise stated, behavior experiments were performed at 25°C and 60% humidity. We performed our locomotion assays in an arena based on previously established FlyBowl^25^ with several modifications. Briefly, a circular piece of white delrin plastic was cut to feature sloped walls according to the FlyBowl dimensions^25^ to construct the arena. A plexiglass cover was cut to serve as the ceiling of the chamber. A custom-built circular LED array featuring IR and red (650nm) lights (LEDSupply) was positioned underneath the arena. Red LEDs were wired via a 1000mA BuckPuck driver (LEDSupply) to enable variable intensities of light. A diffuser sheet was placed above the LED array to ensure lighting was even throughout the arena. LED systems and diffusers were placed inside 3D printed opaque cylinders to focus the LEDs’ light to the arena. The LED system was connected to a Pololu server controller via a relay module to achieve computer control. Fixed above the arena, we positioned a digital camera (FLIR Blackfly S U3-13Y3M-C) with a varifocal lens (LMVZ990-IR) that was fitted with a near-IR bandpass (Midopt BP850) to record the behavior trials. Videos were recorded via Bonsai at 1280×1040 resolution and 30 FPS. All behavior trials were recorded in this arena. All trials except optogenetic activation assays were performed under white light (∼45 μW/cm2) with a polarizing filter to act as a celestial landmark.

### Thermogenetic silencing

Flies that were assayed for our shibire^TS^ experiments were analyzed at either 21°C or 31°C by adjusting the temperature in our behavior chamber. Flies that were analyzed as part of these experiments were placed in 21°C temperature- and humidity-controlled incubators upon eclosion. Flies in the 31°C groups were placed in the chamber for at least 30 minutes before being assayed to allow flies to acclimate to the temperature increase. Flies were analyzed at age 6-9 days old. One fly was subjected to the paradigm at a time. During the assay, we placed a custom-built arrangement of 15 8×8 green LED arrays (Adafruit) on top of the chamber such that the LEDs encircled the behavior arena. LED arrays were wired in parallel and connected to an Arduino to control the position of vertical bar stimulus. Custom Arduino scripts were written for the different stimulus paradigms: clockwise, counterclockwise, and still green vertical bars. Using bonsai, we tracked the fly’s position to calculate its translational velocity in real time. When this value was greater than 1 pixel, a signal is sent to the VR Arduino to permit moving the stimulus. For clockwise and counterclockwise moving vertical bars, the bar positions would update every 25ms–thus a full rotation around the circumference of the arena takes 3s. For the still bars paradigm, one of the eight possible bar positions was chosen at random during the program’s onset and the bar remained still at that position for the duration of the experiment. All 15 8×8 arrays were wired in such a way that each array received the same instructions and thus depicted bars in the same position at each timestep to thus display a uniform visual field.

### Transient optogenetic activation assay

Progeny from crosses for optogenetic activation experiments were divided into two groups. The first group was raised on a diet of standard *Drosophila* media with all-trans retinal (ATR) (Sigma R2500) mixed in with a final concentration of 400 μM. This group is denoted as ATR+. The second group was raised on a diet of standard *Drosophila* media that was mixed with 100% ethanol, the solvent that was used for the ATR. The volume of ethanol added was equivalent to the volume of ATR added in the ATR+ vials. This group is denoted as ATR-. Flies used for optogenetics experiments were raised in an incubator in the absence of any light. Each trial consists of a single male fly. Red light intensity was calibrated to ∼5.0 mW/cm^2^ and shown continuously throughout the stimulus bouts. A single stimulus bout was defined as a single 20s delivery of red light with 20s rest periods before and after stimulus delivery. A small red light was placed in view of the camera during all trials to indicate when the optogenetic stimulus was delivered in each video.

### Sparse activation of columnar neurons

Sparse activation of PFNs was achieved through the previously established SPARC method^30^. PFN SPARC experiments were performed using the nSyb-IVS-PhiC31 SPARC configuration to achieve sparse labeling since the driver line contained off-target neurons that would be more likely to be included in sparse labeling experiments if the UAS-IVS-Phic31 allele was used. We used the UAS-SPARC2-S-CsChrimson::tdTomato allele to achieve sparse labeling since it labeled the smallest proportion of neurons from the starter population. SPARC animals were dissected less than 24 hours after behavioral profiling and then subjected to our IHC protocol. Each fly was stained independently and labeled with a unique identifier number to ensure that each dissected brain preparation could be matched to their respective behavior trials. Trials from 72 total flies were included as part of our final dataset.

### Immunohistochemistry and tissue processing

Immunohistochemistry was performed on adult brains as previously described^47^ but with slight variations. Briefly, adult flies were cold anesthetized on ice and dissected in cold 0.05% PBS-T (T stands for Triton X-100; Fisher Bioreagents, BP151-500). All following steps were performed while brains were nutating. Brains were fixed in 2% PFA/0.5% PBS-T at 4°C overnight. Samples were then washed 4X in 0.5% PBS-T for 15 min each at RT. Brains were then blocked for 30 min at RT in 5% heat-inactivated equine serum (diluted from 100% with 0.5% PBS-T) and then incubated with primary antibodies for two overnights at 4°C. Brains were then again washed 4X in 0.5% PBS-T for 15 min each and then incubated with secondary antibodies for two overnights at 4°C. The samples were then washed again 4X in 0.5% PBS-T for 15 min each before being mounting on a slide (Fisherbrand Superfrost Plus, 12-550-15) in Fluoromount-G mounting medium (SouthernBiotech, 0100-01). The primary antibodies that were used in this study were: Goat anti-GFP (Rockland #600-101-215, 1:1000), Guinea Pig anti-RFP (Gift from Susan Brenner-Morton, Columbia University, 1:10,000), and anti-Brp mouse (nc82, DSHB, 1:50). Secondary antibodies were diluted to 1:1,000. The secondary antibodies used in this study were: donkey anti-Goat Alexa Fluor 488, donkey anti-Guinea pig Alexa Fluor 555, and donkey anti-Mouse 647. Images were taken using confocal microscopy (Zeiss, LSM800) using Zen software (Zeiss). Images were formatted and processed using FIJI (http://fiji.sc).

### Quantification and statistical analysis

#### Analysis of locomotor behavior videos

Automated kinematic tracking of behavior trials was performed via FlyTracker^26^. FlyTracker outputs x and y coordinates and orientation angle for every frame of the behavior videos. Custom R scripts were written to calculate translational and angular velocity from FlyTracker outputs. Translational velocity was calculated as the Euclidean distance between x and y coordinates between successive frames. Angular velocity was calculated as the distance between orientation angle between successive frames. Data was then smoothed using a Gaussian filter with a spread of 10 frames. When computing mean velocity values in our shibire^TS^ experiments, only frames when the fly was less than 30 mm from the center were considered, since this is the region of the of the arena where the flies perform circular walking trajectories. Mean Δ angular velocity and mean Δ translational velocity were calculated as the difference between the means of the velocity value for the frames during which the optogenetic stimulus was off and the frames during which the optogenetic stimulus was on. When translational velocity values were compared in our optogenetics experiments, we normalized translational velocity values by dividing all values by the maximum translational velocity value for that particular fly. This enabled comparison between flies while accounting for variability in each fly’s walking speed. All statistical comparisons were performed using a one-way anova followed by a *post hoc* Tukey test for multiple comparisons. R code that was used to generate figures is available upon request.

#### Quantification of CsChrimson expression

CsChrimson expression was quantified via fluorescence intensity in FIJI. In our PFN SPARC experiments, we determined the level of CsChrimson expression in a nodulus by manually selecting an ROI of a collapsed z-stack of the noduli. We calculated the intensity of labeling as the raw integrated density of signal divided by the area of the noduli. ΔNO_a_ and ΔNO_d_ indices were calculated with the following formula:

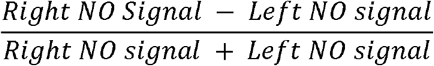

The ΔNO_a-d_ was calculated via the following formula:

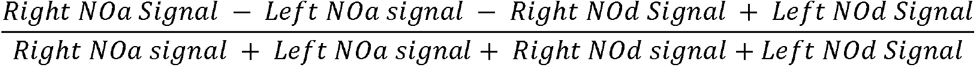

We computed our weighted ΔNO_a-d_ index via the following formula.

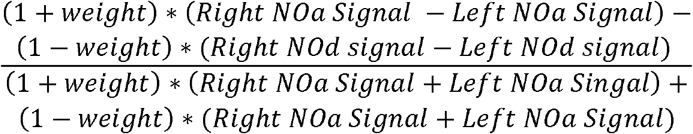

With *weight* being a value that ranges from −1 to 1.

#### ODE model for goal-oriented steering

The system of equations that we used for our model for central complex-mediated steering control in which the goal coordinate was fixed was as follows:

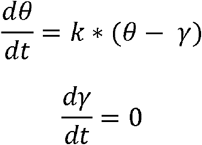

Here, θ equals the fly’s heading direction and γ equals an external goal direction. The constant k represents the value of the fly’s alignment speed, i.e., the rate at which the fly turns to align its heading and goal parameters. For our experiments, we set this constant k to an arbitrary value of 1. Our system of equations solves to the following functions for θ and γ, given the initial values θ_0_ and γ_0_:

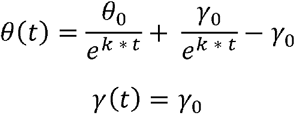

We initialized the values of θ_0_ as 0 and varied γ_0_ between −π and π. We then used these equations to determine the values of dθ/dt and θ from t=0 to t=5. We transformed these values of θ into x and y coordinates in space using the following equations:

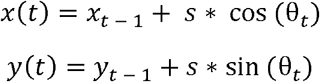

Here, s denotes an arbitrary constant for the fly’s magnitude of translational motion. In our calculations, we used a value of 1 for s. We initialized x_0_ and y_0_ to both equal 0.

In the case where the heading and goal coordinates were constitutively offset, we used the following system of equations:

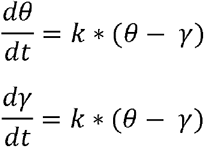

We solved this system with k equaling 1 as above and given the initial values of θ and γ as θ_0_ and γ_0_ respectively to produces the following equations:

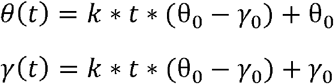

We then set θ_0_ to equal 0 and varied γ_0_ between −π and π as above. We modeled walking trajectories that would arise from the modeled θ values by calculating x, y coordinates for each timepoint as described above.

### Circuit reconstructions of electron microscopy data

Reconstructions of central complex neuropil segmentations and PFN, hDelta, and PFL3 neurons from were accessed via the publicly available Neuprint server for querying data from the hemibrain connectome ^48,49^. 3-D reconstructions were obtained as .obj files and visualized/rendered in blender. Colors were manually selected to correspond to their anatomy. Information on the synaptic connectivity for individual neurons was accessed via Flywire (flywire.ai)^31^. As part of our analysis, we only considered neurons that connect via at least 5 synapses to be connected.

## Data availability

Upon acceptance and publication of this manuscript, all kinematic data and analysis code used for this study will be made available at github.com/anthonycrown/

## Acknowledgements

We thank Dr. Vanessa Ruta, Dr. Charlie Dowell, Dr. Alexander Fleischmann, and the members of the Barnea Laboratory for critical reading of the manuscript. We are grateful to Susan Morton for sharing reagents. We thank John Murphy from the Brown BioMed Machine Shop for aiding in the construction of our behavior chamber. This work was supported by NIH NINDS grant 5R01NS133238 (G.B.), and NIH/NIDCD award F31DC019540 (A.M.C.). Stocks obtained from the Bloomington *Drosophila* Stock Center (NIH P40OD018537) were used in this study.

## Author contributions

A.M.C. and G.B. conceived the project. Behavioral experiments and pre-processing of the resultant data were performed by A.M.C., H.W., and L.H. Immunohistochemical analyses were performed and analyzed by A.M.C. Data analysis and modeling were performed by A.M.C. Data were interpreted by A.M.C. and G.B. The manuscript was written by A.M.C. and G.B.

**Extended Data Table 1.**
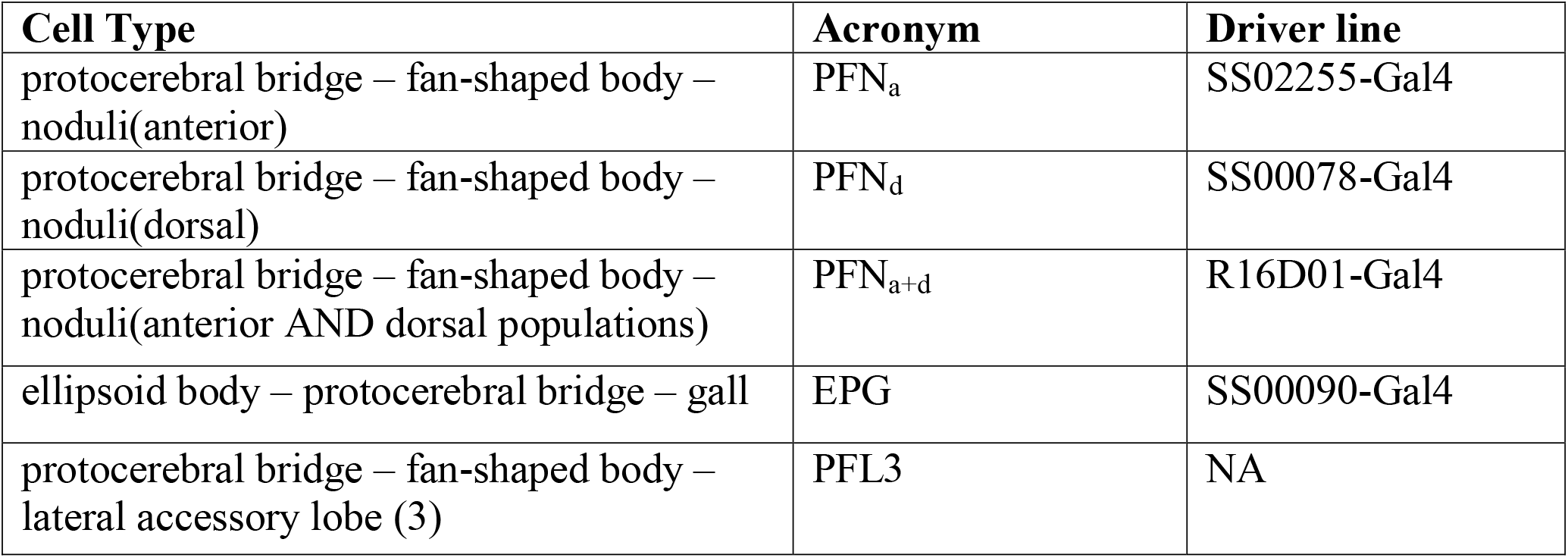
Acronyms used for central complex columnar neurons.

| **Cell Type** | **Acronym** | **Driver line** |
| --- | --- | --- |
| protocerebral bridge – fan-shaped body – noduli(anterior) | PFN<sub>a</sub> | SS02255-Gal4 |
| protocerebral bridge – fan-shaped body – noduli(dorsal) | PFN<sub>d</sub> | SS00078-Gal4 |
| protocerebral bridge – fan-shaped body – noduli(anterior AND dorsal populations) | PFN<sub>a+d</sub> | R16D01-Gal4 |
| ellipsoid body – protocerebral bridge – gall | EPG | SS00090-Gal4 |
| protocerebral bridge – fan-shaped body – lateral accessory lobe (3) | PFL3 | NA |

**Extended Data Table 2.** Genotypes used for each figure.

| **Figure** | **Shorthand** | **Genotype** |
| --- | --- | --- |
| Fig. 1f-m | PFNd >Shi<sup>TS</sup> | w<sup>1118</sup>; R16D01-p65ADZp/+ ; R15E01-ZpGdbd/UAS-Shi<sup>TS</sup> |
| Fig. 1l, m | PFNa >Shi<sup>TS</sup> | w<sup>1118</sup>; R16D01-p65ADZp/+ ; VT016114-ZpGdbd/UAS-Shi<sup>TS</sup> |
| Fig. 1l, m, Extended Data Fig. 2 | EPG >Shi<sup>TS</sup> | w<sup>1118</sup>; R19G12-p65ADZp/+ ; R15C03-ZpGdbd/UAS-Shi<sup>TS</sup> |
| Fig. 1l, m, Extended Data Fig. 2 | No driver > Shi<sup>TS</sup> | w<sup>1118</sup>; +/+ ; UAS- Shi<sup>TS</sup>/+ |
| Fig. 2e-h, Extended Data Fig. 4 | No driver > CsChrimson | w<sup>1118</sup>; UAS-CsChrimson ; +/+ |
| Fig. 2e-h, Extended Data Fig. 4 | PFNa > CsChrimson | w<sup>1118</sup>; R16D01-p65ADZp/ UAS-CsChrimson ; R15E01-ZpGdbd/+ |
| Fig. 2e-h, Extended Data Fig. 4 | PFNd > CsChrimson | w<sup>1118</sup>; R16D01-p65ADZp/ UAS-CsChrimson ; VT016114-ZpGdbd/+ |
| Fig. 2e-h, Extended Data Fig. 4, 5 | PFNa+d > CsChrimson | w<sup>1118</sup>; UAS-CsChrimson ; R16D01-Gal4/+ |
| Fig. 3 | R16D01-Gal4 > SPARC | nSyb-IVS-PhiC31 ; UAS-SPARC2-S-CsChrimson::TdTomato/+; R16D01-Gal4 |
| Fig. 3i-l, Extended Data Fig. 6 | SS00081 > CsChrimson | w<sup>1118</sup>; R16D01-p65ADZp/ UAS-CsChrimson ; R21H11-ZpGdbd/+ |
| Extended Data Fig. 2 | WT | w<sup>1118</sup>;; |

**Extended Data Figure 1.**
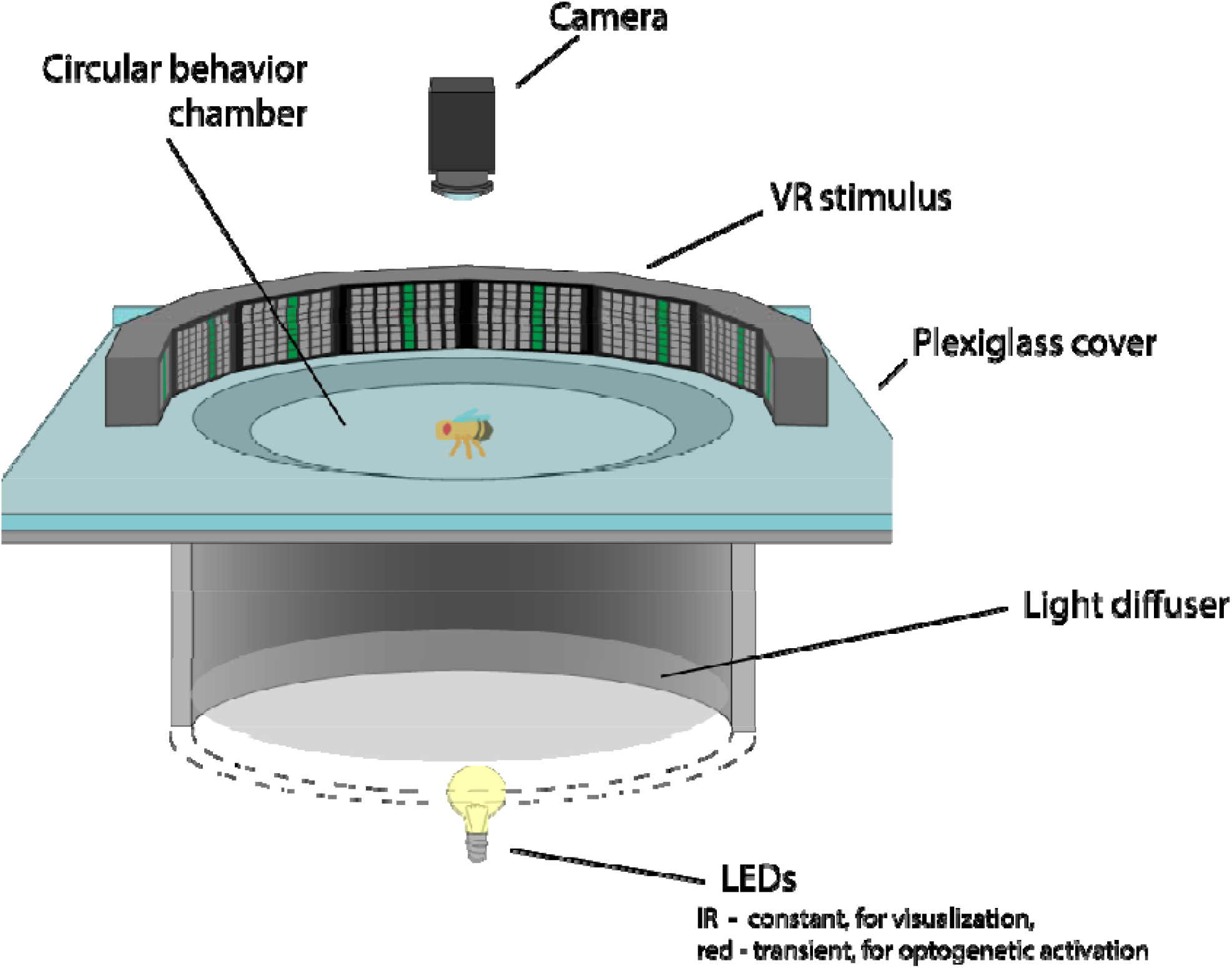
Apparatus used for quantifying locomotor behaviors. Schematic of the custom-made behavior arena used for the locomotor tracking experiments. Cross-section of VR stimulus and lighting chamber is shown. Flies are placed in the arena beneath a plexiglass ceiling while a camera records their behavior.

**Extended Data Figure 2.**
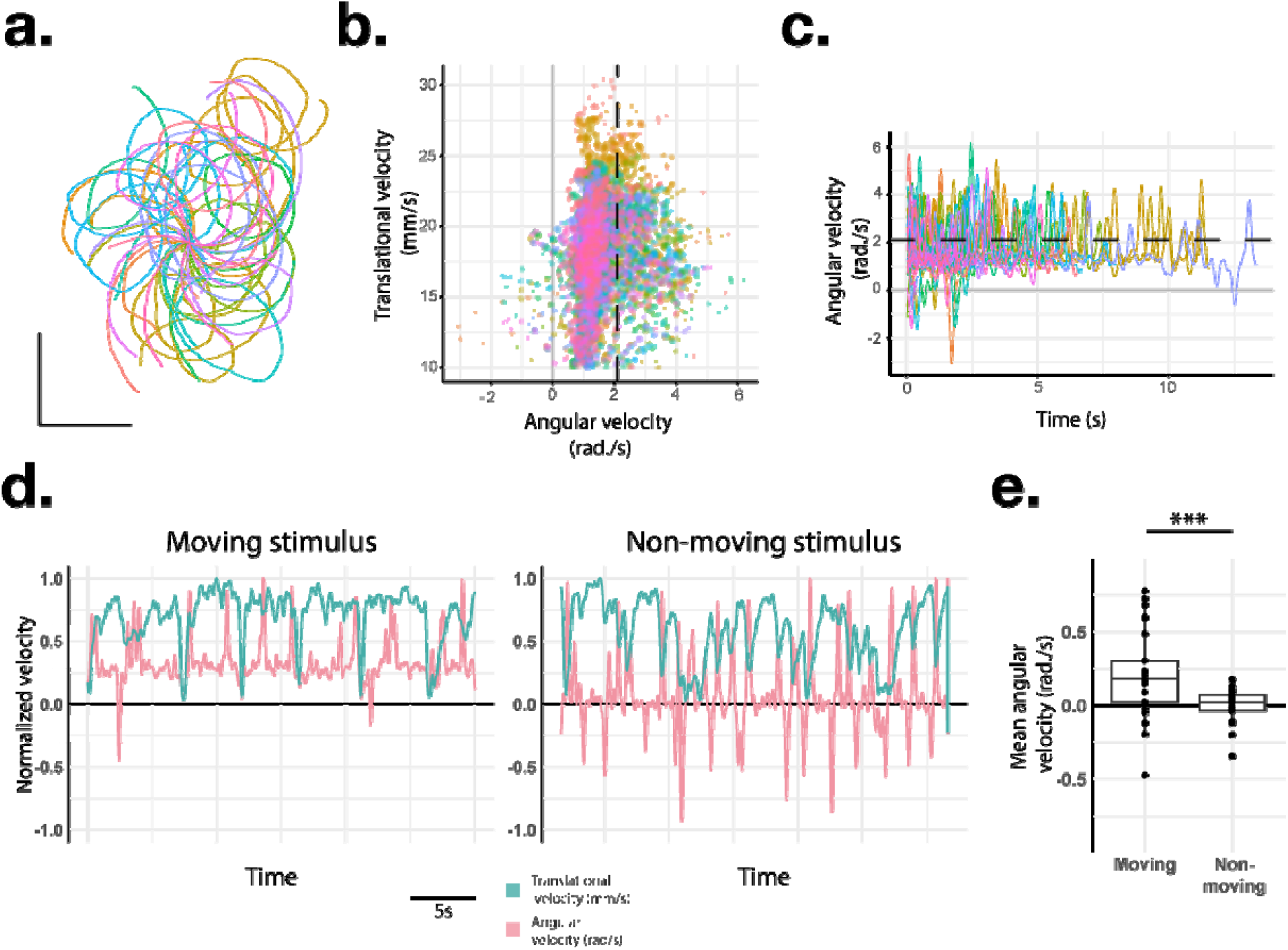
Quantifying reorientation bouts in WT flies. **(a-c)** Representative data from angular motion behavioral paradigm. One trial from a WT fly is depicted. Each color depicts a single walking bout. (a) walking trajectories whose x and y coordinates were translated to a common starting position. (b) scatter plot of the translational and angular velocity values for each timepoint. (c) line plot of angular velocity values over time. Dashed line in (b, c) depicts rotational velocity of VR stimulus. **(d)** Line plots depicting translational and angular velocity values for a single fly during 30s of walking in the VR stimulus paradigm. Translational and angular velocity values are coincident when the VR stimulus bar is moving (left panel) but not when the bar is still (right panel). Velocity values are normalized such that the maxima equal 1 to enable comparisons between translational and angular velocity values. **(e)** boxplot of angular velocity values Average of entire 5 min trial is shown. Positive angular velocity values indicate towards the direction of the stimulus. Scale bar in (a) equals 10mm. Wilcoxon ranked sum test was used for statistical testing (*** = p < 0.001)

**Extended Data Figure 3.**
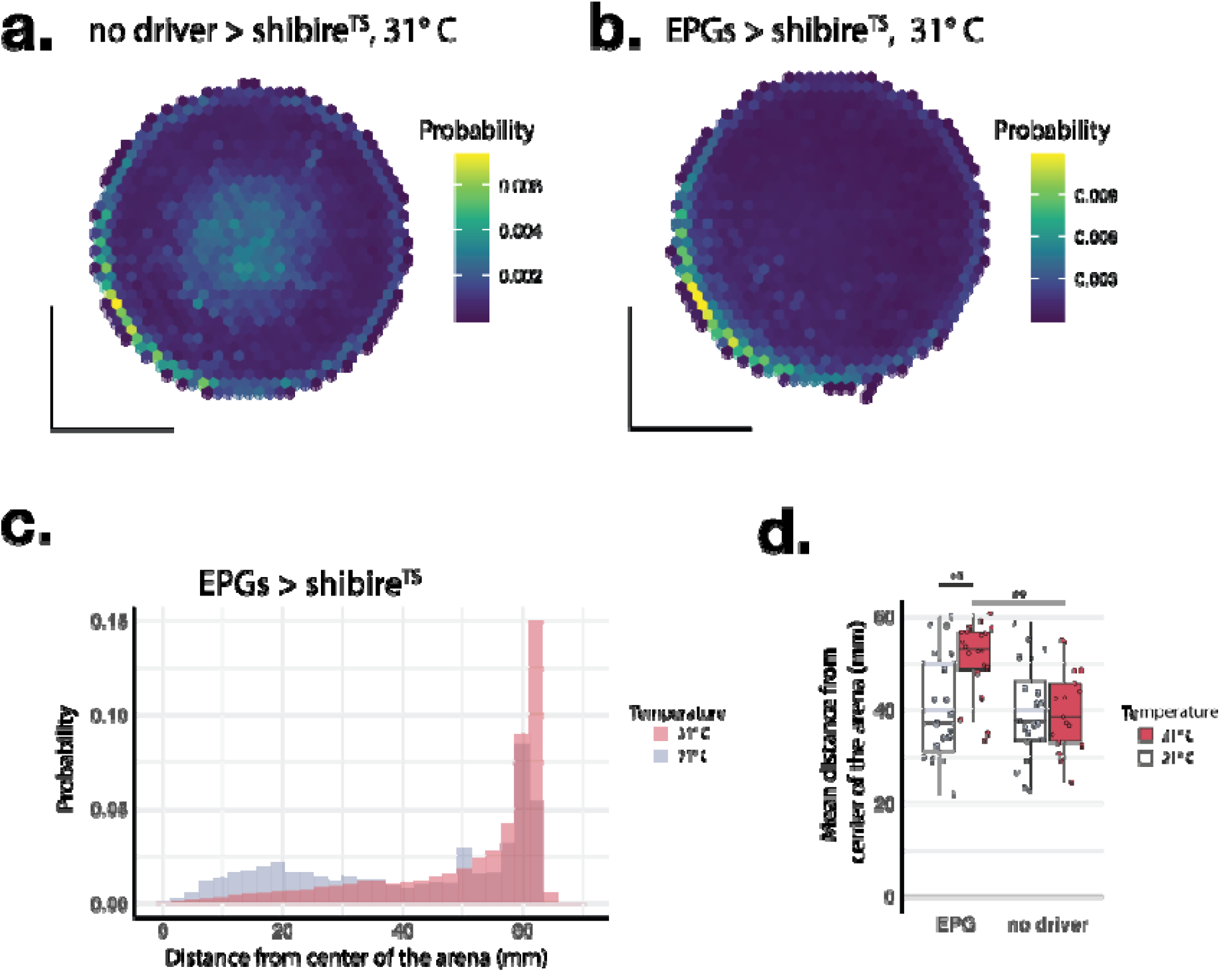
Reorientation bouts require neurotransmission from EPGs. **(a, b)** Two-dimensional density plots depicting the probability of flies’ x/y coordinates across trials during our optic flow assay. (a) depicts the 31° C trials with control flies carrying shibire^TS^ but no genetic driver. (b) depicts the 31° C trials where flies expressed shibire^TS^ in EPG neurons. When EPG neurons were silenced by shibire^TS^, flies were noticeably less present in the center of the arena, the region where we observe reorientation bouts. **(c)** Probability histogram of the distance from the center of the arena for each time point across trials for the EPG > shibire^TS^ group. 21° C (grey) and 31° C (red) trials are plotted. Note that values are bimodal in the 21° C condition, with one peak centered close to the center of the arena and another centered at the edge while values in the 31° C condition are unimodal with one peak at the edge of the arena. **(d)** Boxplot of values of mean distances from the center of the behavior arena. Silencing EPGs led to flies walking predominantly at the edges of the arena, while we observe no such trend in the other groups.

**Extended Data Figure 4.**
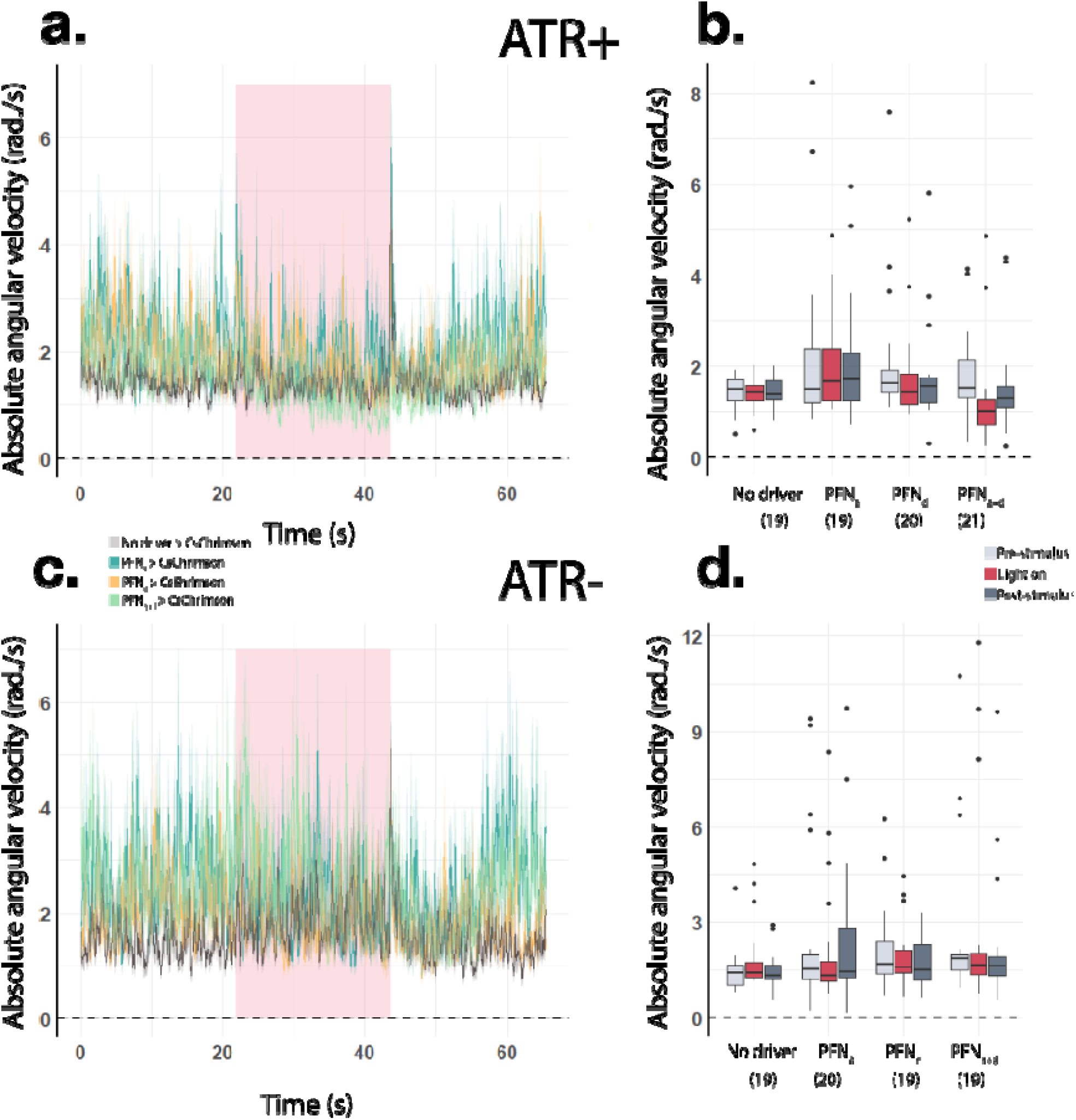
Angular velocity values do not change during bulk activation of PFNs. **(a-d)** Quantification of angular velocity values from optogenetic experiments depicted in Fig. 2e-h. (a) Line plot of mean absolute angular velocity values (±s.e.) across stimulus bouts for the various genotypes in the ATR+ condition. Red box denotes time interval of delivery of the optogenetic stimulus. See (b) for the numbers of trials (N) that were averaged in each group. (b) Boxplot of mean absolute angular velocity values during each trial in the ATR+ condition. Values are shown for the optogenetic stimulus period as well as for the pre- and post-stimulus periods. (c, d) Same as (a, b) but for the ATR-groups.

**Extended Data Figure 5.**
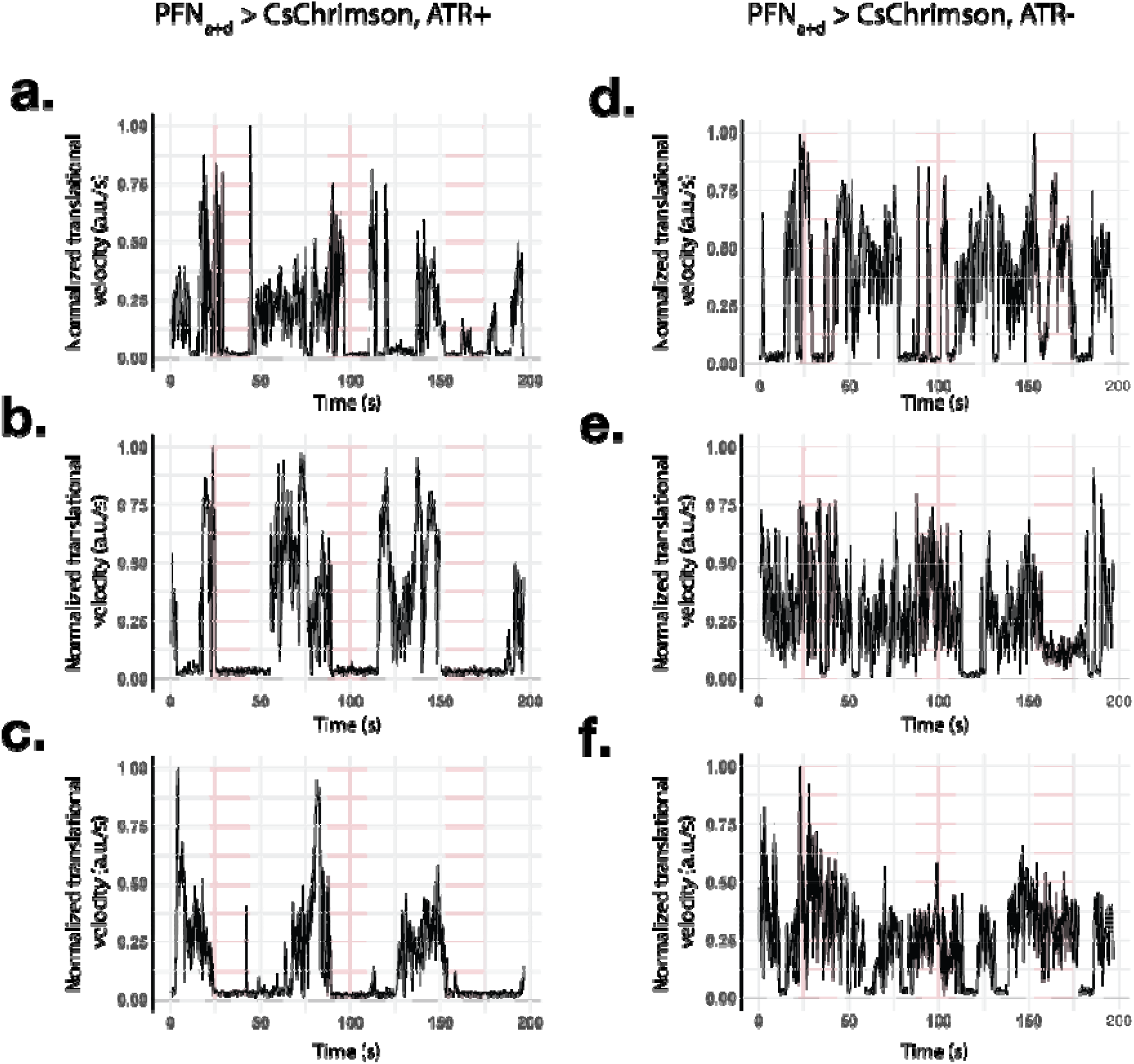
Representative trials for freezing behavior during PFN_a+d_ activation. **(a-c)** Normalized translational velocity values plotted over time for PFN_a+d_ activation trials for the ATR+ condition. Each plot depicts normalized translational velocity values for an individual fly over the course of an experiment. Red boxes indicate the time interval in which the optogenetic stimulus was delivered. **(d-f)** Same as (a-c) but in the ATR-condition. Translational velocity values drop to zero during optogenetic stimulus delivery in the ATR+ condition but not the ATR-condition.

**Extended Data Figure 6.**
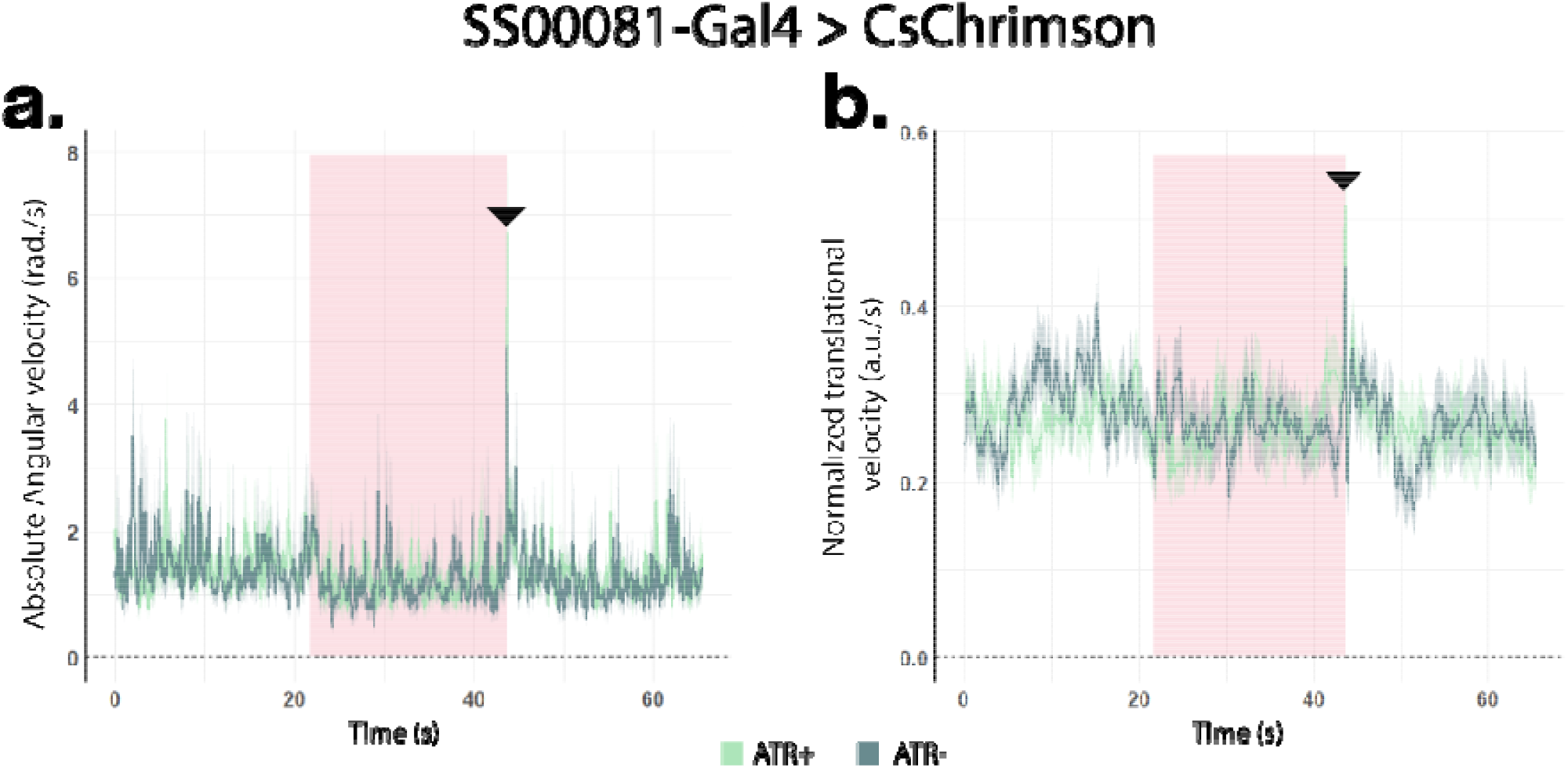
Sparse symmetric activation of PFNs elicits no change in locomotion. **(a, b)** Line plots of averaged absolute angular velocity (a) and normalized translational velocity (b) values (±s.e.) while optogenetically activating the sparse but symmetric population of PFNs that is targeted by SS00081-Gal4. Trials where flies were raised on a diet supplemented with ATR (ATR+) (N=10) or without (ATR-) (N=11) are shown. Red box indicates time interval of delivery of optogenetic stimulus. Spikes in angular and translational velocity are observed upon offset of the optogenetic stimulus in both ATR+ and ATR-conditions (black arrowheads).

## Notes

### Competing Interest Statement

The authors have declared no competing interest.

## Bibliography

1 Müller, M. & Wehner, R. Vol. 94 589–594 (Naturwissenschaften, 2007).

2 Merlin, C. & Liedvogel, M. Vol. 222 (J of Exp Biol., 2019).

3 Heinze, S. & Reppert, S. M. Vol. 69 345–358 (Neuron, 2011).

4 Honkanen, A., Adden, A., Da Silva Freitas, J. & Heinze, S. Vol. 222 (J. Exp. Biology, 2019).

5 Nguyen, T. A. T., Beetz, M. J., Merlin, C., El Jundi, B. & Vol. 288 (Proc. R. Soc. B., 2021).

6 Hulse, B. K. et al. Vol. 10 (eLife, 2021).

7 Wolff, T. & Rubin, G. M. Vol. 526 2585–2611 (J Comp Neurol, 2018).

8 Kim, S. S., Hermundstad, A. M., Romani, S., Abbott, L. F. & Jayaraman, V. Vol. 576 126–131 (Nature, 2019).

9 Shiozaki, H. M., Ohta, K. & Kazama, H. Vol. 106 126-141.e125 (Neuron, 2020).

10 Turner-Evans, D. et al. Vol. 6 (eLife, 2017).

11 Giraldo, Y. M. et al. Vol. 28 2845-2852.e2844 (Curr Biol, 2018).

12 Okubo, T. S., Patella, P., Alessandro, I. D. A., Wilson, R. I. & Vol. 107 924-940.e918 (Neuron, 2020).

13 Dan, C., Hulse, B. K., Kappagantula, R., Jayaraman, V. & Hermundstad, A. M. A neural circuit architecture for rapid learning in goal-directed navigation. Neuron (2024). 10.1016/j.neuron.2024.04.036

14 Lu, J. et al. Vol. 601 98–104 (Nature, 2022).

15 Currier, T. A., Matheson, A. M. M. & Nagel, K. I. Vol. 9 1–29 (eLife, 2020).

16 Lyu, C., Abbott, L. F. & Maimon, G. Vol. 601 92–97 (Nature, 2022).

17 Matheson, A. M. M. et al. Vol. 13 1–21 (Nat Commun, 2022).

18 Hu, W. et al. Vol. 24 1573–1584 (Cell Rep, 2018).

19 Sareen, P. F., McCurdy, L. Y. & Nitabach, M. N. Vol. 12 (Nat Commun, 2021).

20 Goldschmidt, D. et al. 2023.2007.2019.549514 (bioRxiv, 2023).

21 Mussells Pires, P., Zhang, L., Parache, V., Abbott, L. F. & Maimon, G. Converting an allocentric goal into an egocentric steering signal. Nature 626, 808–818 (2024). 10.1038/s41586-023-07006-3

22 Westeinde, E. A. et al. Transforming a head direction signal into a goal-oriented steering command. Nature 626, 819–826 (2024). 10.1038/s41586-024-07039-2

23 Warren, T. L., Weir, P. T. & Dickinson, M. H. Vol. 221 (J Exp Biol., 2018).

24 Green, J., Vijayan, V., Mussells Pires, P., Adachi, A. & Maimon, G. A neural heading estimate is compared with an internal goal to guide oriented navigation. Nat Neurosci 22, 1460–1468 (2019). 10.1038/s41593-019-0444-x

25 Simon, J. C. & Dickinson, M. H. Vol. 5 (ed Kenji Hashimoto) e8793 (PLoS One, 2010).

26 Eyjolfsdottir, E. et al. Vol. 8690 772–787 (Computer Vision — ECCV 2014, 2014).

27 Green, J. et al. Vol. 546 101–106 (Nature, 2017).

28 Seelig, J. D. & Jayaraman, V. Vol. 521 186–191 (Nature, 2015).

29 Yamaguchi, S., Desplan, C. & Heisenberg, M. Vol. 107 5634–5639 (Proc Natl Acad Sci U S A, 2010).

30 Isaacman-Beck, J. et al. Vol. 23 1168–1175 (Nat Neurosci, 2020).

31 Dorkenwald, S. et al. Neuronal wiring diagram of an adult brain. bioRxiv (2023). 10.1101/2023.06.27.546656

32 Bidaye, S. S. et al. Vol. 108 469-485.e468 (Neuron, 2020).

33 Fujiwara, T., Brotas, M. & Chiappe, M. E. Walking strides direct rapid and flexible recruitment of visual circuits for course control in Drosophila. Neuron 110, 2124–2138 e2128 (2022). 10.1016/j.neuron.2022.04.008

34 Chiappe, M. E. Circuits for self-motion estimation and walking control in Drosophila. Curr Opin Neurobiol 81, 102748 (2023). 10.1016/j.conb.2023.102748

35 Cruz, T. L., Perez, S. M. & Chiappe, M. E. Fast tuning of posture control by visual feedback underlies gaze stabilization in walking Drosophila. Curr Biol 31, 4596–4607 e4595 (2021). 10.1016/j.cub.2021.08.041

36 Kim, I. S. & Dickinson, M. H. Idiothetic Path Integration in the Fruit Fly Drosophila melanogaster. Curr Biol 27, 2227–2238 e2223 (2017). 10.1016/j.cub.2017.06.026

37 Corfas, R. A., Sharma, T. & Dickinson, M. H. Vol. 29 1660-1668.e1664 (Cell Press, 2019).

38 Behbahani, A. H., Palmer, E. H., Corfas, R. A. & Dickinson, M. H. Drosophila re-zero their path integrator at the center of a fictive food patch. Curr Biol 31, 4534–4546 e4535 (2021). 10.1016/j.cub.2021.08.006

39 Ishida, I. G., Sethi, S., Mohren, T. L., Abbott, L. F. & Maimon, G. Neuronal calcium spikes enable vector inversion in the Drosophila brain. bioRxiv (2023). 10.1101/2023.11.24.568537

40 Bicanski, A. & Burgess, N. Vol. 21 453–470 (Nat Rev Neurosci, 2020).

41 Taube, J. S., Muller, R. U. & Ranck, J. B., Jr. Head-direction cells recorded from the postsubiculum in freely moving rats. II. Effects of environmental manipulations. J Neurosci 10, 436–447 (1990). 10.1523/jneurosci.10-02-00436.1990

42 Taube, J. S., Muller, R. U. & Ranck, J.B., Jr. Head-direction cells recorded from the postsubiculum in freely moving rats. I. Description and quantitative analysis. J Neurosci 10, 420–435 (1990). 10.1523/jneurosci.10-02-00420.1990

43 Taube, J. S. The head direction signal: origins and sensory-motor integration. Annu Rev Neurosci 30, 181–207 (2007). 10.1146/annurev.neuro.29.051605.112854

44 Hartley, T., Burgess, N., Lever, C., Cacucci, F. & O’Keefe, J. Modeling place fields in terms of the cortical inputs to the hippocampus. Hippocampus 10, 369–379 (2000). 10.1002/1098-1063(2000)10:4<369::Aid-hipo3>3.0.Co;2-0

45 Grieves, R. M., Duvelle, É. & Dudchenko, P. A. A boundary vector cell model of place field repetition. Spatial Cognition & Computation 18, 217–256 (2018). 10.1080/13875868.2018.1437621

46 Kitamoto, T. Vol. 47 81–92 (J Neurobiol, 2001).

47 Talay, M. et al. Vol. 96 783-795.e784 (Neuron, 2017).

48 Bates, A., Jefferis, G. & Franconville, R. (https://natverse.org/neuprintr, https://github.com/natverse/neuprintr., 2021).

49 Scheffer, L. K. et al. Vol. 9 1–74 (eLife, 2020).

